# Spike-specific CXCR3^+^ T_FH_ cells play a dominant functional role in supporting antibody responses in SARS-CoV-2 infection and vaccination

**DOI:** 10.1101/2022.08.09.503302

**Authors:** Jian Zhang, Rongzhang He, Bo Liu, Xingyu Zheng, Qian Wu, Bo Wen, Qijie Wang, Ziyan Liu, Fangfang Chang, Yabin Hu, Ting Xie, Yongchen Liu, Jun Chen, Jing Yang, Shishan Teng, Tingting Bai, Yanxi Peng, Ze Liu, Yuan Peng, Weijin Huang, Velislava Terzieva, Youchun Wang, Wenpei Liu, Yi-Ping Li, Xiaowang Qu

## Abstract

CD4^+^ T follicular helper (T_FH_) cells are required for high-quality antibody generation and maintenance. However, the longevity and functional role of these cells are poorly defined in COVID-19 convalescents and vaccine recipients. Here, we longitudinally investigated the dynamics and functional roles of spike-specific circulating T_FH_ cells and their subsets in convalescents at the 2^nd^, 5^th^, 8^th^, 12^th^ and 24^th^ months after COVID-19 symptom onset and in vaccinees after two and three doses of inactivated vaccine. SARS-CoV-2 infection elicited robust spike-specific T_FH_ cell and antibody responses, of which spike-specific CXCR3^+^ T_FH_ cells but not spike-specific CXCR3^−^ T_FH_ cells and neutralizing antibodies were persistent for at least two years in more than 80% of convalescents who experienced symptomatic COVID-19, which was well coordinated between spike-specific T_FH_ cell and antibody responses at the 5^th^ month after infection. Inactivated vaccine immunization also induced spike-specific T_FH_ cell and antibody responses; however, these responses rapidly declined after six months with a two-dose standard administration, and a third dose significantly promoted antibody maturation and potency. Functionally, spike-specific CXCR3^+^ T_FH_ cells exhibited better responsiveness than spike-specific CXCR3^−^ T_FH_ cells upon spike protein stimulation in vitro and showed superior capacity in supporting spike-specific antibody secreting cell (ASC) differentiation and antibody production than spike-specific CXCR3^−^ T_FH_ cells cocultured with autologous memory B cells. In conclusion, spike-specific CXCR3^+^ T_FH_ cells played a dominant functional role in antibody elicitation and maintenance in SARS-CoV-2 infection and vaccination, suggesting that induction of CXCR3-biased spike-specific T_FH_ cell differentiation will benefit SARS-CoV-2 vaccine development aiming to induce long-term protective immune memory.

**Highlights:**

- SARS-CoV-2 infection elicited robust spike-specific T_FH_ cell and antibody responses, which persisted for at least two years in the majority of symptomatic COVID-19 convalescent patients.
- Inactivated vaccine immunization also elicited spike-specific T_FH_ cell and antibody responses, which rapidly declined over time, and a third dose significantly promoted antibody maturation and potency.
- Spike-specific CXCR3^+^ T_FH_ cells exhibited more durable responses than spike-specific CXCR3^−^ T_FH_ cells, correlated with antibody responses and showed superior capacity in supporting ASC differentiation and antibody production than spike-specific CXCR3^−^ T_FH_ cells.

## Introduction

Severe acute respiratory syndrome coronavirus 2 (SARS-CoV-2) is the causative agent of coronavirus disease 2019 (COVID-19), which has posed a serious health threat and considerable socioeconomic consequences to the world population(Zhou et al., 2020; Zhu et al., 2020). Effective strategies are urgently needed to establish persistent and appropriate magnitude immune memory at the individual and population levels to prevent the continued spread of infection. Thus, understanding the successful immune memory in COVID-19-recovered and vaccinated individuals is essential for vaccine design to elicit long-lasting humoral and cellular immune responses against SARS-CoV-2 and variants of concern (VOCs).

Both SARS-CoV-2 natural infection and vaccination have been reported to be able to elicit robust humoral and cellular immune responses(Long et al., 2020; Meckiff et al., 2020; Mudd et al., 2022; Painter et al., 2021; Peng et al., 2020; Rydyznski Moderbacher et al., 2020; Sekine et al., 2020; Thevarajan et al., 2020; Weiskopf et al., 2020). Although immune memory has also been shown to demonstrate relatively stable B and T-cell memory in the observation period, the antibody levels are found to be waning out over time(Ibarrondo et al., 2020; Long *et al*., 2020; Roltgen and Boyd, 2021). The early appearance of neutralizing antibodies (nAbs) associated with less severe disease in acute COVID-19 and the persistence of nAbs in recovered individuals contribute to preventing reinfection by blocking virus entry(Dupont et al., 2021; Zhou et al., 2021). To date, it is believed that reinfection of endemic human coronaviruses (HCoV-229E, HCoV-OC43, HCoV-NL63, and HCoV-HKU1) occurs frequently, primarily due to the short-lived immunoglobins induced by the endemic coronaviruses(Edridge et al., 2020). In contrast, three highly pathogenic coronaviruses (SARS-CoV-1, MERS-CoV, and SARS-CoV-2) trigger full immune defences, eliciting stronger adaptive immunity in most recovered cases, although accumulating reinfection cases with VOCs have been reported(Cao et al., 2007; Cheon et al., 2022; Yang et al., 2022). Persistent and appropriate levels of nAbs could block virus entry, thus preventing the reinfection cycle of SARS-CoV-2 or emerging VOCs(Altmann and Boyton, 2021).

The production and maturation of high-affinity antibodies and memory B cells and long-lived plasma cell differentiation mainly rely on germinal centre (GC) reactions in secondary lymphoid tissues, which are tightly regulated by T follicular helper (T_FH_) cells(Crotty, 2019). T_FH_ cells are specialized B helper cells that enable the proliferation, survival, and differentiation of GC B cells through the delivery of costimulatory molecules and cytokine signals(Johnston et al., 2009; Nurieva et al., 2009; Schaerli et al., 2000; Yu et al., 2009). Circulating T_FH_ cells could serve as counterparts of GC T_FH_ cells, as they express low levels of PD-1, ICOS, and Bcl6 and exhibit a memory phenotype(Morita et al., 2011). These characteristics have been correlated with high-affinity antibody responses during virus infection and vaccination(Bentebibel et al., 2016; Bentebibel et al., 2013; Martin-Gayo et al., 2017; Niessl et al., 2020; Zhang et al., 2019).

In the acute phase of SARS-CoV-2 infection, large amounts but low-affinity antibodies were produced rapidly, and in parallel, antigen-specific CD4^+^ T cells and circulating T_FH_ cells appeared, which in turn contributed to antibody production to combat infection(Chen et al., 2020; Le Bert et al., 2020; Meckiff *et al*., 2020; Zhou *et al*., 2021). In COVID-19 convalescents, we and others have previously demonstrated that CXCR3^+^ T_FH_ cells directly correlate with anti-spike antibody responses(Gong et al., 2020; Juno et al., 2020; Zhang et al., 2021). Several studies also show that CCR6^+^ T_FH_ cells (most are CXCR3^−^ T_FH_ cells) were dominant and could be maintained longer after recovery in the observation period(Dan et al., 2021; Juno *et al*., 2020; Rodda et al., 2021; Rydyznski Moderbacher *et al*., 2020). However, the longevity and functional role of these cells in antibody elicitation and maintenance are not clear. Vaccination with the mRNA vaccine elicited strong as well as COVID-19 convalescent-specific humoral immunity, including circulating T_FH_ cell responses(Painter *et al*., 2021; Sahin et al., 2021). More recently, several studies have shown that SARS-CoV-2 infection and mRNA vaccination efficiently induce robust GC reactions, GC-resident T_FH_ and B-cell responses, and long-lived plasma cells in bone marrow(Kim et al., 2022; Laidlaw and Ellebedy, 2022; Lederer et al., 2022; Turner et al., 2021a; Turner et al., 2021b). These responses ensure antibody production to prevent reinfection or maintain the level of circulating antibody. Although severely impaired GC reactions were found in critically ill patients or dying patients, the majority of infected or vaccinated individuals were reported to have GC reactions in lymphoid tissues for several months, which may support the maturation and maintenance of high-affinity antibodies(Duan et al., 2020; Kaneko et al., 2020; Laidlaw and Ellebedy, 2022; Poon et al., 2021; Turner *et al*., 2021a). Nevertheless, addressing the longevity of antibody and T_FH_ cell responses as well as the functional role of T_FH_ cells in supporting the antibody response in COVID-19 convalescents and vaccinated subjects is urgently required to guide vaccine development.

In this study, we longitudinally investigated the kinetics and longevity of spike-specific antibody and T_FH_ cell responses in SARS-CoV-2 infection and vaccination and addressed the functional roles of spike-specific circulating T_FH_ cells and their subsets in supporting ASC differentiation and antibody production. Our findings provide new insights into the development of interventions and vaccines against SARS-CoV-2 and VOCs.

## Results

### Persistence of spike-specific circulating T_FH_ cell responses in COVID-19 convalescents for at least 12 months

To longitudinally assess circulating T_FH_ cell and antibody responses after recovery from COVID-19, 104 blood samples were collected from 37 convalescents at the 2^nd^, 5^th^, 8^th^, 12^th^, and 24^th^ months after COVID-19 symptom onset (Figure 1A and Supporting Table 1), and PBMCs were isolated and cultured for 24 h with the stimulation of SARS-CoV-2 spike protein or BSA (5 μg/ml). Negative control PBMCs were collected from 17 healthy individuals before the COVID-19 pandemic and stimulated with the same conditions (Figure. 1B). Circulating T_FH_ (CXCR5^+^ CD4^+^ T) cells were gated, and a CD154 (CD40 L) assay was applied to measure antigen-specific T cells (Supporting Figure 1). The results showed that the frequencies of CD154^+^ T_FH_ cells at the 2^nd^, 5^th^, 8^th^ and 12^th^ months, but not at the 24^th^ month, were significantly higher in the spike-stimulated group than in the BSA group; healthy PMBCs showed no difference in either stimulation (Figure 1B-C). These results suggest that SARS-CoV-2 infection induced spike-specific T_FH_ cell responses, which could last for at least 12 months after recovery. We also analysed spike-specific non-T_FH_ (CXCR5^−^ CD4^+^ T) cell responses and showed that these spike-responsive cells exhibited a similar pattern to that of T_FH_ cells maintained for at least 12 months (Supporting Figure 2A). Longitudinal analysis revealed that the frequencies of spike-specific T_FH_ cells markedly declined from the 2^nd^ to the 5^th^ month and then remained at a similar level for up to 24 months. The responsiveness capacity presented as the stimulation index (SI: spike-responsive versus BSA-responsive [baseline]) was relatively stable at each time point (Figure 1D). The kinetics of spike-specific non-T_FH_ cells resembled those of spike-specific T_FH_ cells; however, the spike-specific non-T_FH_ cell responsiveness capacity declined significantly from the 2^nd^ to the 5^th^ months (SI of the 2^nd^ month and later time points; Supporting Figure 2B). These findings suggest that SARS-CoV-2 infection elicited robust spike-specific T_FH_ cell and non-T_FH_ cell responses, of which responsiveness could be maintained for at least one year.

**Figure 1.**
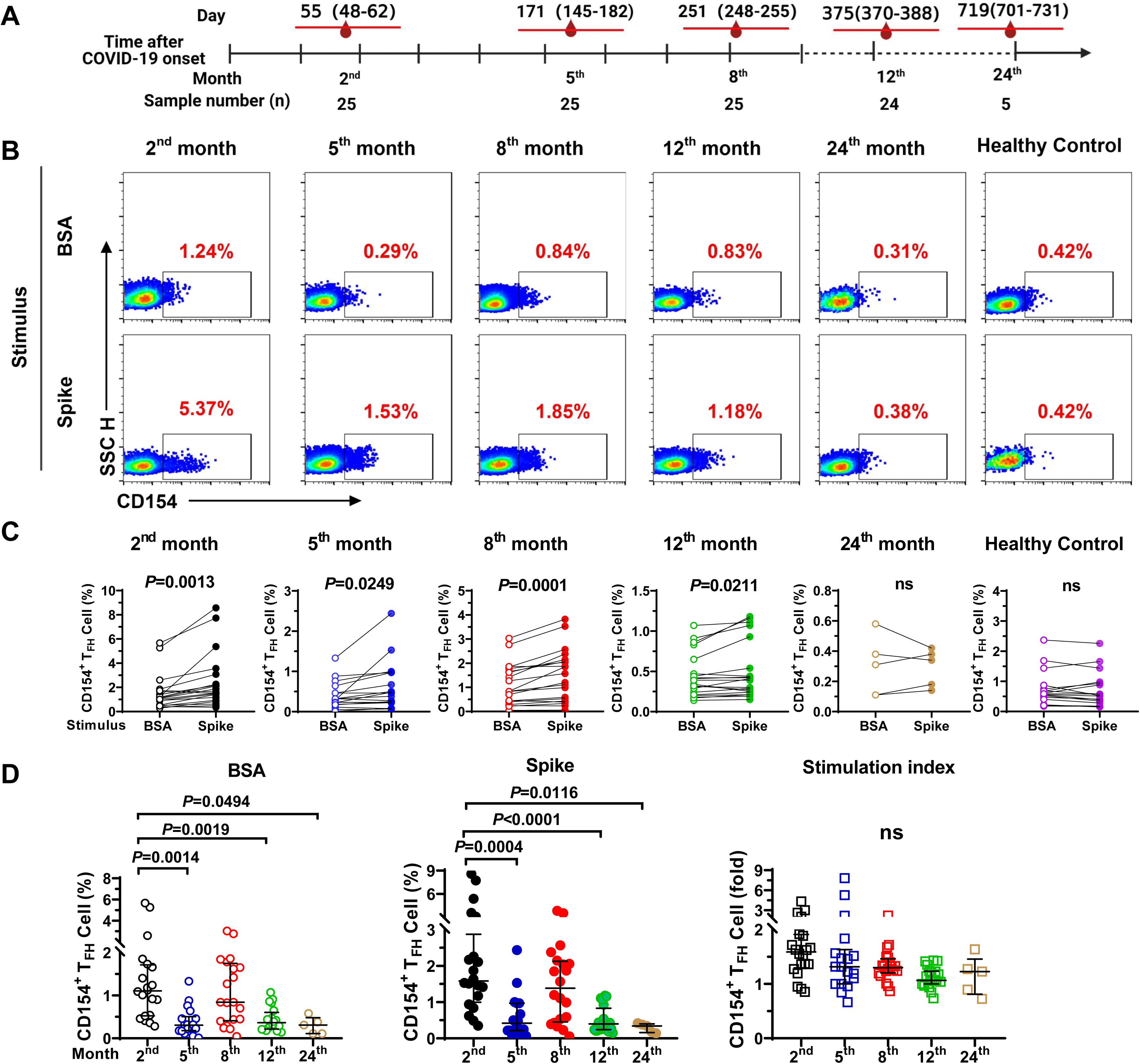
Longitudinal analysis of T_FH_ cell responses in COVID-19 convalescents. (A) Diagram of sample collection from COVID-19 convalescents. The median (interquartile range, IQR) of days of sampling and the number of samples are shown for each time point. PBMCs were isolated from each blood sample for later analysis. (B) Representative flow plots of spike-specific T_FH_ cells (CD154^+^) upon BSA or spike protein stimulation. (C) Spike-specific T_FH_ cells at the 2^nd^ month (n=20), 5^th^ month (n=18), 8^th^ month (n=20), 12^th^ month (n=20), and 24^th^ month (n=5), as well as healthy controls (n=17). A paired t test was used to analyse the difference between BSA and spike protein stimulation. (D) Kinetics of spike-specific T_FH_ cell responses and the stimulation index (SI). One-way analysis of variance was used to compare the differences among multiple groups, and Tukey’s multiple-comparisons test was used to compare differences within the groups. The data are presented as the median ±LJIQR (25%–75%). *P*<LJ0.05 was considered to be a two-tailed significant difference, ns means not significant.

### Spike-specific but distinct recall responses of CXCR3^+^ and CXCR3^−^ T_FH_ cells in COVID-19 convalescents

Previously, cross-section studies showed that the frequencies of CXCR3^+^ T_FH_ cells or T_H_1-like T_FH_ (T_FH_1) cells were correlated with spike-specific antibody responses in COVID-19 convalescents(Boppana et al., 2021; Gong *et al*., 2020; Juno *et al*., 2020; Zhang *et al*., 2021). To explore the kinetics and longevity of T_FH_ cell subset responses (Supporting Figure 1), stimulated PBMCs were further gated into CXCR3^+^ and CXCR3^−^ T_FH_ cell subsets, followed by analysis of CD154 expression. The results showed that CD154^+^ CXCR3^+^ T_FH_ cells from the 2^nd^ to the 24^th^ month were significantly enhanced after spike stimulation compared with BSA stimulation, while spike-responsive CXCR3^−^ T_FH_ cells were only observed for those from the 2^nd^ to the 8^th^ month, and no significant changes were observed for the cells from the 12^th^ to the 24^th^ month (Figure 2A-B). Longitudinally, both the frequencies of spike-specific CXCR3^+^ T_FH_ cells and spike-specific CXCR3^−^ T_FH_ cells declined from the 2^nd^ to the 5^th^ month but remained relatively durable from the 5^th^ to the 24^th^ month (Figure 2C). The SIs of spike-specific CXCR3^+^ and CXCR3^−^ T_FH_ cells did not significantly vary throughout 24 months (Figure 2C). Of note, the proportion of spike-specific CXCR3^−^ T_FH_ cells was higher than that of spike-specific CXCR3^+^ T_FH_ cells at each time point, but spike-specific CXCR3^+^ T_FH_ cells exhibited superior responsiveness than spike-specific CXCR3^−^ T_FH_ cells (higher SIs) at each time point upon stimulation (Figure 2D). PD-1 expression represents the active form of circulating T_FH_ cells, and it was found that PD-1^+^ T_FH_ cells gradually declined over time after the resolution of COVID-19(Dan *et al*., 2021). Here, we found that the frequencies of spike-specific PD-1^+^ CXCR3^+^ T_FH_ cells from the 2^nd^ to the 12^th^ month were significantly higher in the spike-stimulated group than in the BSA group, whereas the responsiveness of spike-specific PD-1^+^ CXCR3^−^ T_FH_ cells was only found at the 2^nd^ month (Supporting Figure 3A-B), although spike-specific PD-1^+^ CXCR3^−^ T_FH_ cells had relatively higher frequencies than spike-specific PD-1^+^ CXCR3^+^ T_FH_ cells (Supporting Figure 3C). In addition, spike-responsive CXCR3^+^ non-T_FH_ cells were maintained up to 12 months, while spike-specific CXCR3^−^ non-T_FH_ cell responses were only observed to the 8^th^ month upon spike stimulation (Supporting Figure 4A-B). The kinetic patterns of both spike-specific CXCR3^+^ and CXCR3^−^ non-T_FH_ cells were similar to those of spike-specific CXCR3^+^ and CXCR3^−^ T_FH_ cells, respectively (Supporting Figure 4C). The SIs of spike-specific CXCR3^+^ non-T_FH_ cells were higher than those of CXCR3^−^ non-T_FH_ cells from the 2^nd^ to the 12^th^ months (Supporting Figure 4D). Furthermore, spike-specific T_FH_ and non-T_FH_ cells, spike-specific CXCR3^+^ T_FH_ and CXCR3^+^ non-T_FH_ cells, and spike-specific CXCR3^−^ and CXCR3^−^ non-T_FH_ cells were highly correlated, respectively, from the 2^nd^ to the 12^th^ month, except at the 24^th^ month, due to a limited sample size (Supporting Figure 5A-C), indicating that well-balanced spike-specific T_FH_ cell and non-T_FH_ cell responses were elicited in SARS-CoV-2 infection. Together, these data demonstrated that spike-specific CXCR3^+^ and CXCR3^−^ T_FH_ cells exhibited distinct recall responses, and spike-specific CXCR3^+^ T_FH_ cells maintained a longer active state and better responsiveness than spike-specific CXCR3^−^ T_FH_ cells in COVID-19 convalescents.

**Figure 2.**
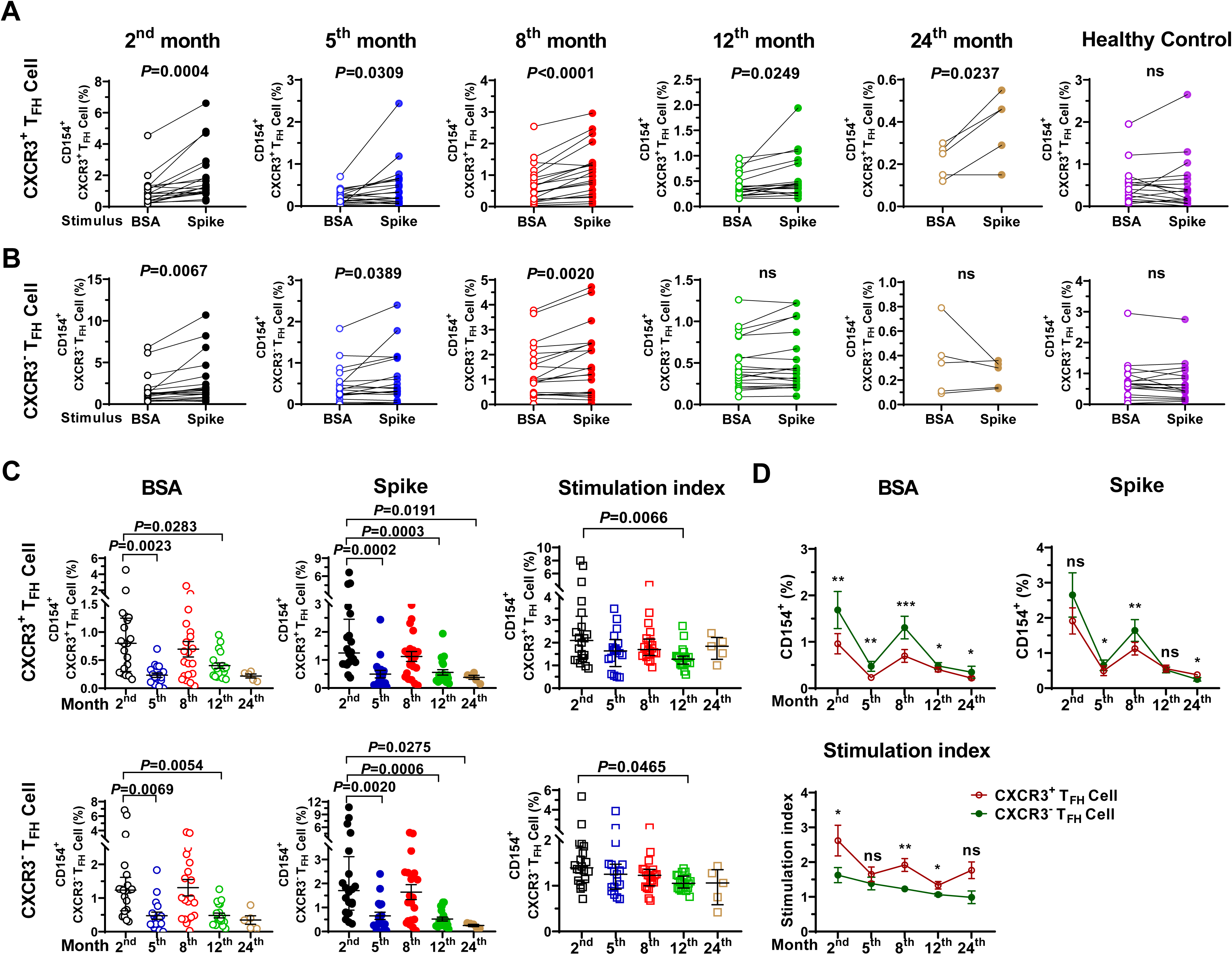
Distinct responses of spike-specific CXCR3^+^ and CXCR3^−^ T_FH_ cell subsets. (A-B) Spike-specific CXCR3^+^ and CXCR3^−^ T_FH_ cell responses at the 2^nd^ month (n=20), 5^th^ month (n=18), 8^th^ month (n=20), 12^th^ month (n=20) and 24^th^ month (n=5) upon BSA or spike protein stimulation. A paired t test was used to analyse the difference in BSA and spike protein stimulation. (C) Kinetics of spike-specific CXCR3^+^ and CXCR3^−^ T_FH_ cell responses and stimulation index (SI). One-way analysis of variance was used to compare the differences among multiple groups, and Tukey’s multiple-comparisons test was used to compare differences within the groups. Data are the median ± IQR (25%– 75%). (D) Comparison of spike-specific CXCR3^+^ and CXCR3^−^ T_FH_ cell frequency and SI. A paired t test was used to analyse the difference between CXCR3^+^ and CXCR3^−^ T_FH_ cell frequency or SI with BSA and spike protein stimulation. * *P*<0.05; ** *P*<0.01; *** *P*<0.001; **** *P*<0.0001. *P*< 0.05 was considered to be a two-tailed significant difference, ns means not significant.

### Dynamics of spike-specific antibody responses in COVID-19 convalescents and correlations with T_FH_ cells

Next, to assess the kinetics of antibody responses in COVID-19 convalescents, we examined spike-specific antibodies in plasma at the 2^nd^, 5^th^, 8^th^, 12^th^ and 24^th^ months. The endpoint titer and avidity of spike-specific antibodies were determined, such as immunoglobin A (IgA), IgG and subclasses (IgG1, IgG2, and IgG3). Endpoint titers of spike-specific IgG, IgG1, IgG3, and IgA were detectable at each time point, but all declined significantly from the 2^nd^ to the 5^th^ month and then remained stable from the 5^th^ to the 24^th^ month (Figure 3A). However, spike-specific IgG2 was at a low level or undetectable at each time point (data not shown). IgA has been reported to rapidly decline, and most are short-lived(Sterlin et al., 2021). Here, IgA was found to persist for at least 24 months in symptomatic convalescents. In contrast, the avidity of the spike-specific IgG, IgG1, IgG3, and IgA antibodies increased over time with different kinetics from endpoint titers, indicating that the antibody continued to mature during the convalescent phase (Figure 3B). This finding was in line with the observation in recovered SARS patients, in which antibody avidity continued to mature from low avidity in early phase to high avidity at late phase of disease recovery(Chan et al., 2005). The maturation of antibody avidity also reflected the persistence of GC reactions after SARS-CoV-2 resolution(Poon *et al*., 2021).

**Figure 3.**
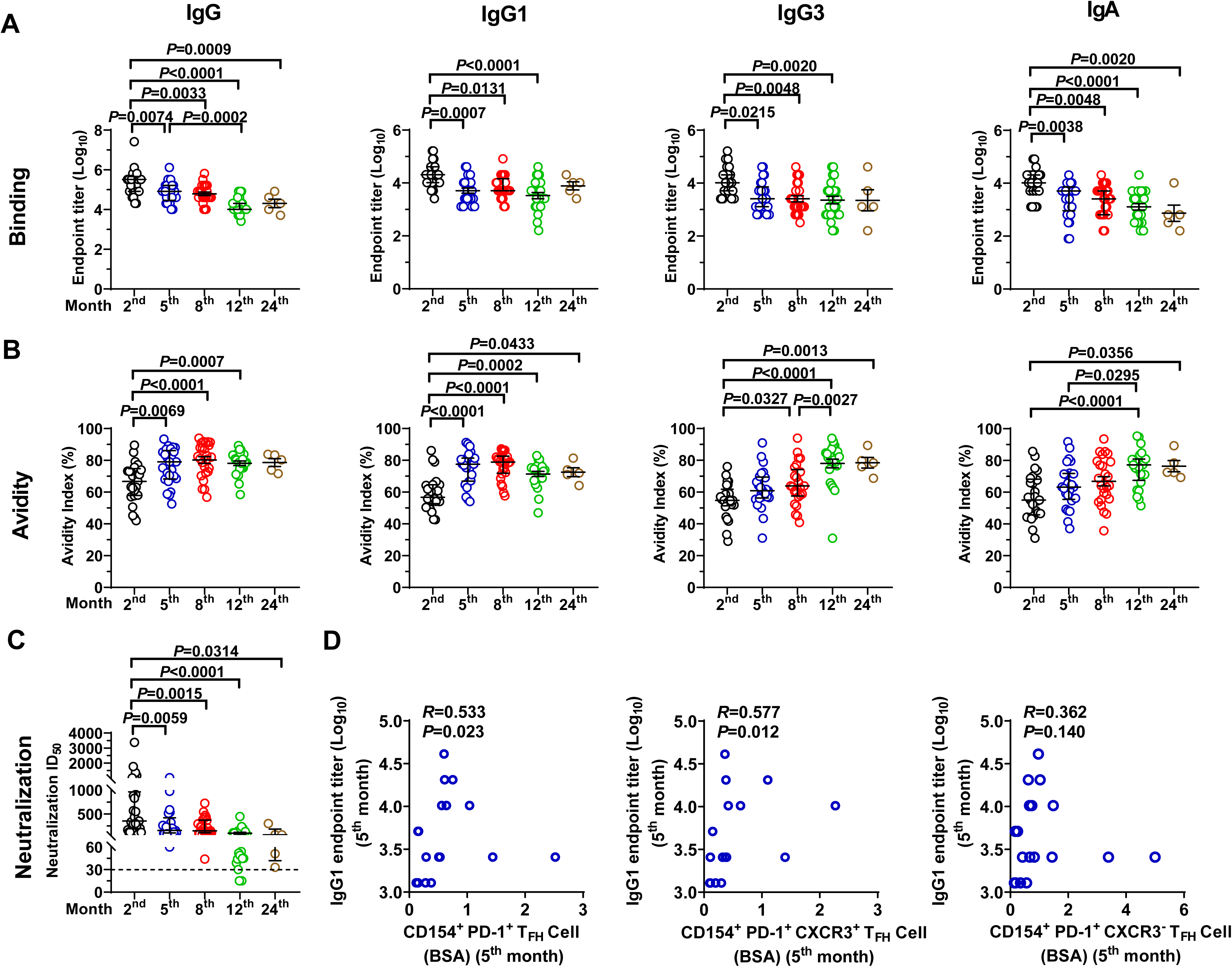
Kinetics of spike-specific antibody responses in COVID-19 convalescents. (A) Endpoint titers and (B) avidity index of plasma spike-specific IgG, IgG1 IgG3, and IgA antibodies in COVID-19 convalescent patients at the 2^nd^ month (n=25), 5^th^ month (n=25), 8^th^ month (n=25), 12^th^ month (n=24), and 24^th^ month (n=5) after illness onset. The endpoint titer and avidity index data were logarithmically transformed. (C) Neutralization titers of COVID-19 convalescent against SARS-CoV-2 pseudotyped virus at the indicated time points. In A, B, and C, one-way analysis of variance was used to compare the differences among multiple groups, and Tukey’s multiple-comparisons test was used to compare differences within groups. Data are presented as the median ± IQR (25%–75%). (D) Correlations of spike-specific PD-1^+^ T_FH_, spike-specific PD-1^+^ CXCR3^+^ and PD-1^+^ CXCR3^−^ T_FH_ cells with IgG1 endpoint titers at the 5^th^ month. Spearman’s rank correlation coefficient was used to describe the association between the frequencies of T_FH_ cells and subsets with the IgG1 endpoint titers. *P*<0.05 was considered a significant difference in a two-sided test.

Then, we examined the neutralization activity of a plasma antibody against SARS-CoV-2 spike pseudotyped virus. The neutralization titers significantly decreased from the 2^nd^ to the 5^th^ month and then gradually declined from the 5^th^ to the 24^th^ month (Figure 3C). Within the first 8 months, all individuals showed neutralizing antibody titers above the cut-off value (≥30); however, the titers declined, and 83% (20 out of 24) and 80% (4 out of 5) were maintained until the 12^th^ and 24^th^ months, respectively (Figure 3C). These data suggest that the neutralizing antibody dropped in the early phase of recovery, but the majority was maintained at a lower level for at least two years, which has similar patterns to the dynamics of the antibody endpoint titers (Figure 3A). The neutralization titers were positively associated with the endpoint titers of IgG, IgG1, IgG3, and IgA from the 2^nd^ to the 12^th^ month; less significant correlations were found at the 24^th^ month, which included only five samples (Supporting Figure 6A-D). These results suggest that both IgG and IgA titers were persistent and contributed to the neutralization effect for at least two years in the majority of convalescents.

The dynamics of spike-specific T_FH_ cells and their subsets and antibody titers all declined but shared a similar pattern from the 5^th^ to the 24^th^ month (Figure 1–2, Figure 3A and 3C). Thus, the correlations of spike-specific T_FH_ cells or their subsets and antibody titers at each time point were further analysed. The endpoint titers of IgG1 were positively correlated with the frequencies of CD154^+^ PD-1^+^ T_FH_ and CD154^+^ PD-1^+^ CXCR3^+^ T_FH_ cells but not CD154^+^ PD-1^+^ CXCR3^−^ T_FH_ cells, and notably, these correlations were only observed at the 5^th^ month (Figure 3D). No correlations were found between the IgG1 antibody titer and resting T_FH_ cells or their subset or non-T_FH_ cells or their subset at each time point (Supporting Figure 7A-F). Together, these results further demonstrated the potential role of spike-specific CXCR3^+^ T_FH_ cells in assisting antibody elicitation and maintenance in COVID-19.

### Longitudinal analysis of spike-specific T_FH_ cell and antibody responses in inactivated vaccine recipients

SARS-CoV-2 infection elicited robust and highly correlated antibody and T_FH_ cell responses, and such responses were maintained for up to two years. To assess the dynamics of neutralizing antibodies and T_FH_ cells following vaccination, we recruited 26 participants who completed the standard vaccination procedure of the two-dose inactivated vaccine and collected blood samples from multiple time points to analyse the antibody titers and T_FH_ cell responses (Figure 4A and Supporting Table 2). The neutralization antibody reached the peak titer 14 days after the second dose of vaccination (45 days after the first dose), and then the titers rapidly decreased to a lower level at Day 180 (Figure 4B). Different from neutralization kinetics, the IgG avidity index exhibited a lower level at Days 14, 28, and 45 and then reached a plateau at Day 90 through Day 180 (Figure 4C). Notably, the peak of the vaccine-derived antibody avidity index was lower than that of natural infection with SARS-CoV-2 at the 8^th^ month (natural infection peak index versus vaccination index: median, IQR: 82.00%, 72.42%–90.21% vs. 55.37%, 49.93%–61.15%). These findings suggest that a two-dose immunization with inactivated vaccine elicited a lower level of neutralizing antibody and that these antibodies were not fully mature (low avidity) compared with the natural infection of SARS-CoV-2.

**Figure 4.**
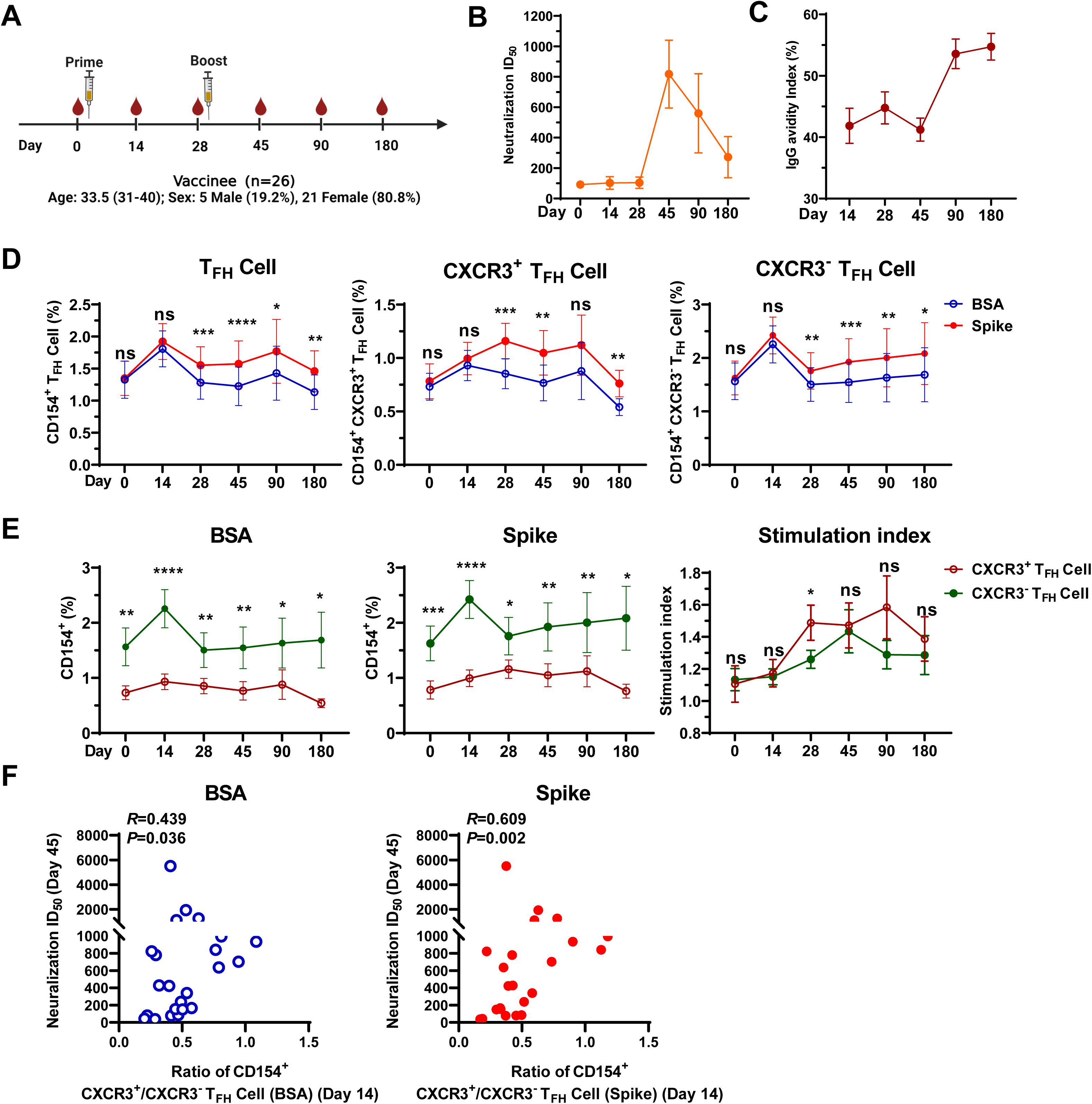
The kinetics of spike-specific antibody and circulating T_FH_ cell responses in inactivated vaccine recipients. (A) Strategic diagram of vaccination and bleeding. (B) Neutralizing antibody titers against pseudotyped SARS-CoV-2 spike virus at the indicated time points ([Day 14, n=26; Day 28, n=26; Day 45, n=26; Day 90, n=25; Day 180, n=25]). Data are the mean ± SEM. (C) The kinetics of spike-specific IgG avidity at different time points ([Day 14, n=26; Day 28, n=26; Day 45, n=26; Day 90, n=25; Day 180, n=25]). Data are the mean ± SEM. (D) Frequencies of spike-specific T_FH_ cells and CXCR3^+^ and CXCR3^−^ T_FH_ cells before (Day 0, n=24) and after vaccination (Day 14, n=24; Day 28, n=23; Day 45, n=24; Day 90, n=26; Day 180, n=19). A paired t test was used to analyse the difference between T_FH_ cells and subsets between BSA and spike protein stimulation at the indicated time points, and the data are the mean ± SEM. (E) Comparison of spike-specific CXCR3^+^ and CXCR3^−^ T_FH_ cell responses as well as the SI at the indicated time points upon BSA or spike protein stimulation. A paired t test was used to analyse the difference between CXCR3^+^ and CXCR3^−^ T_FH_ cells at the indicated time points, and the data are the mean ± SEM. (F) Correlation between the ratio of spike-specific CXCR3^+^/CXCR3^−^ T_FH_ cells on Day 14 with neutralization antibody on Day 45 (n=24). Spearman’s rank coefficient of correlation was used for T_FH_ cell and neutralization antibody analysis. * *P*<0.05; ** *P*<0.01; *** *P*<0.001; **** *P*<0.0001. *P*<0.05 was considered to be a two-tailed significant difference, ns means not significant.

Accordingly, the frequencies of spike-specific T_FH_ and CXCR3^−^ T_FH_ cells peaked at Day 14 after the first dose, followed by a rapid decrease and then a slight elevation after the second dose (Day 28) and were maintained at this level for at least 180 days (Figure 4D). In contrast, spike-specific CXCR3^+^ T_FH_ cells increased slowly and peaked at Day 28, were maintained up to Day 90, and then showed a sharp decrease at Day 180 (Figure 4D). Importantly, spike-specific CXCR3^+^ T_FH_ cells showed stronger responsiveness (higher SIs) than spike-specific CXCR3^−^ T_FH_ cells, although spike-specific CXCR3^−^ T_FH_ cells had higher frequencies than spike-specific CXCR3^+^ T_FH_ cells upon BSA or spike stimulation, which was in line with natural infection (Figure 4E). Both spike-specific PD-1^+^ CXCR3^+^ and PD-1^+^ CXCR3^−^ T_FH_ cells showed similar kinetics, levels, and responsiveness to spike-specific CXCR3^+^ T_FH_ cells and CXCR3^−^ T_FH_ cells, respectively (Supporting Figure 8A-C). No direct correlations between the frequencies of spike-specific PD-1^+^ T_FH_ cells or PD-1^+^ CXCR3^+^ T_FH_ cells and antibody titers were observed which exhibited at the 5^th^ month in natural infection (data not shown). To explore whether there is a link between T_FH_ cells and antibody responses in inactivated vaccine recipients, we analysed the ratio of spike-specific PD-1^+^ CXCR3^+^/PD-1^+^CXCR3^−^ T_FH_ cells and found that the ratio of spike-specific PD-1^+^ CXCR3^+^/PD-1^+^ CXCR3^−^ T_FH_ cells at Day 14 after the first dose were positively correlated with the peak neutralization titers of the second dose (Day 45) (Figure 4F). These data suggest that early primed T_FH_ cells, especially spike-specific CXCR3^+^ T_FH_ cells, contribute to antibody production in the late phase, highlighting a key role of spike-specific CXCR3^+^ T_FH_ cells in the early activation of CD4^+^ T cells. In addition, spike-specific non-T_FH_ cells were also induced by the inactivated vaccine, and spike-specific CXCR3^+^ non-T_FH_ cells were superior in response to stimulation (Supporting Figure 9A-C). These results indicate that two-dose vaccination with inactivated vaccine could efficiently elicit neutralizing antibody and activate T_FH_ cell and non-T_FH_ cell responses; however, the magnitude and persistence of these immune responses were weak and short compared to natural infection.

### A third dose booster augments the T_FH_ cell response and promotes spike-specific antibody potency and maturation

Because the antibody response waned significantly after 6 months with a two-dose standard regimen, a third dose booster was recommended. To test the antibody and T_FH_ cell responses before and after the third-dose vaccination, we recruited 24 individuals who had completed a two-dose regimen for at least 6 months and collected PBMCs before and 14 days after the third dose boost (Supporting Table 2). The third dose dramatically increased the neutralization titer more than 12-fold (Figure 5A) and significantly promoted antibody affinity maturation (Figure 5B), reaching a level similar to that of natural infection (natural infection peak index versus vaccination index: median, IQR: 82.00%, 72.42%–90.21% vs. 86.11%, 84.99%–88.42%). T_FH_ cells and CXCR3^+^ and CXCR3^−^ T_FH_ cell subsets all responded to spike stimulation before and after the third dose booster (Figure 5C); however, the expansion of spike-specific CXCR3^+^ T_FH_ cells was apparently faster and higher than that of spike-specific CXCR3^−^ T_FH_ cells (Figure 5D-E). Furthermore, the ratio of spike-specific PD-1^+^ CXCR3^+^/ PD-1^+^ CXCR3^−^ T_FH_ cells before the booster was significantly correlated with the neutralization titers after the booster (Figure 5F). These results suggested that the spike-specific T_FH_ cells elicited by the two-dose regimen further supported antibody production following the third dose booster. Spike-specific non-T_FH_ cells and CXCR3^+^ and CXCR3^−^ non-T_FH_ cell subsets were also expanded by the third dose (Supporting Figure 10A-B), but spike-specific CXCR3^+^ non-T_FH_ cells were more efficiently expanded than spike-specific CXCR3^−^ non-T_FH_ cells upon spike stimulation (Supporting Figure 10 C). Thus, these data showed that the third dose booster augmented the responses of antibody, spike-specific T_FH_ cells and their subsets and non-T_FH_ cells, of which spike-specific CXCR3^+^ T_FH_ cells were preferentially expanded and contributed to antibody quality and quantity enhancement.

**Figure 5.**
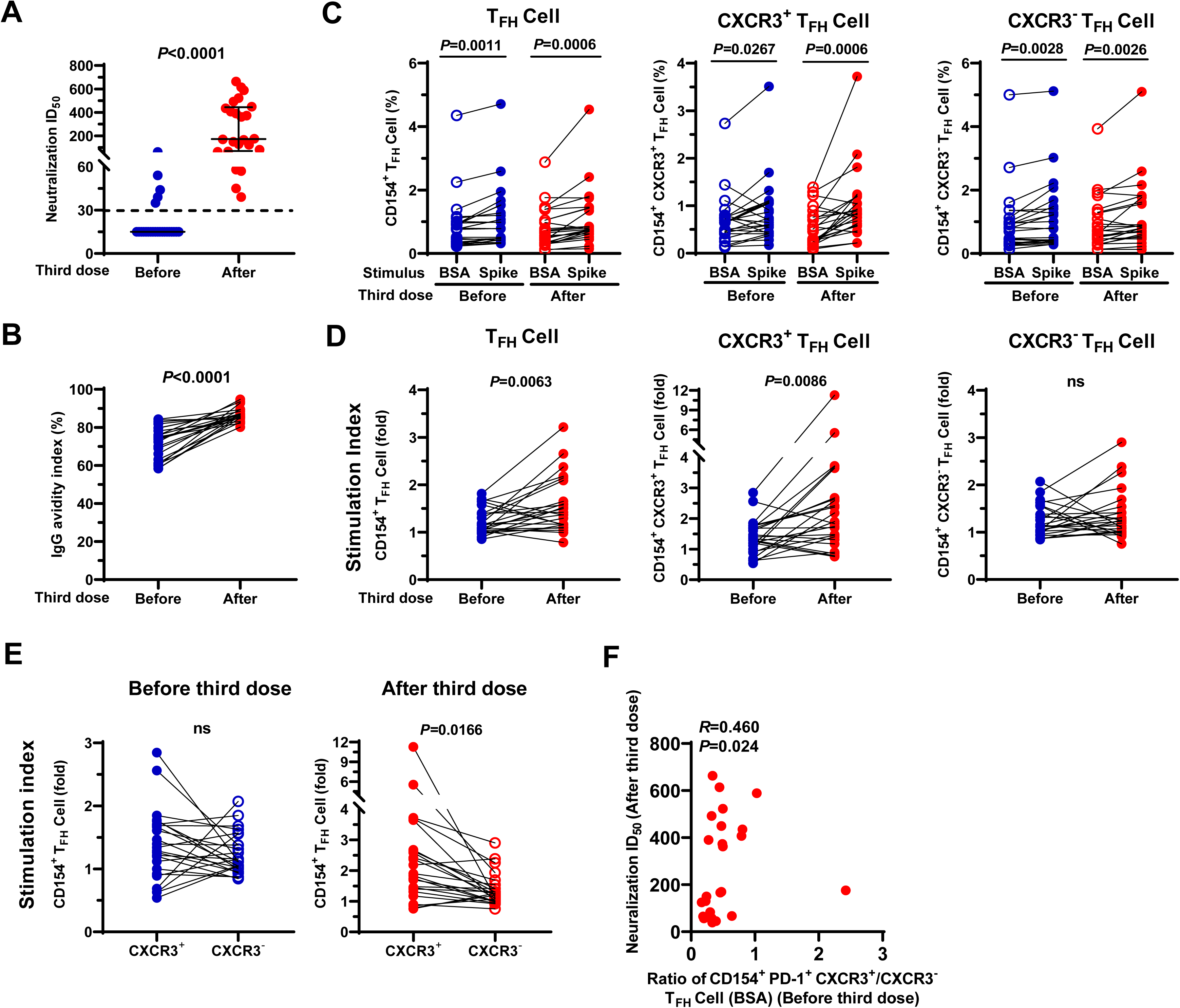
The third dose booster promoted spike-specific antibody production and maturation. (A) Neutralization titers before and after the third dose boost (n=24), Mann Whitney U test was used to compare the difference between two groups, data were median ± IQR (25%–75%). (B) IgG avidity before and after the third dose boost (n=24). (C) Frequencies of spike-specific T_FH_ cells and CXCR3^+^ and CXCR3^−^ T_FH_ cells before and after the third dose boost upon BSA or spike protein stimulation (n=24). (D) The SI of spike-specific T_FH_ cells and CXCR3^+^ and CXCR3^−^ T_FH_ cells before and after the third dose boost (n=24). (E) Comparison of SIs of spike-specific CXCR3^+^ and CXCR3^−^ T_FH_ cells before and after the third dose boost (n=24). In B-E, a paired t test was used to analyse the difference between two groups. (F) Correlation of the ratio of spike-specific PD-1^+^ CXCR3^+^/CXCR3^−^ T_FH_ cells before the third dose and neutralization titers after the third dose boost (n=24). Spearman’s rank coefficient of correlation was used for the correlation of T_FH_ cells and neutralization. *P*<0.05 was considered a significant difference in a two-sided test, ns means not significant.

### Spike-specific CXCR3^+^ T_FH_ cells from natural infection and vaccination show superior capacity than spike-specific CXCR3^−^ T_FH_ cells in supporting antibody-secreting cell differentiation and antibody production in vitro

Previous studies have shown that both spike-specific CD4^+^ T cells and T_FH_ cells are associated with antibody production in SARS-CoV-2 infection(Gong *et al*., 2020; Juno *et al*., 2020; Peng *et al*., 2020; Reynolds et al., 2020; Zhang *et al*., 2021). To discriminate the functional role of these cells in supporting ASC differentiation and antibody production, we cocultured T_FH_ cells or non-T_FH_ cells with autologous memory B cells (5×10^4^ cells for each cell type) (Figure 6A). The T_FH_ cells or non-T_FH_ cells from each group of healthy controls, convalescents, and vaccine recipients were cocultured with autologous memory B cells for 6 days in the presence of *staphylococcal enterotoxin* B (SEB), total ASCs (CD38^hi^ CD27^hi^ B cells) and spike-specific ASCs were measured by FACS (Figure 6B), and spike-specific IgG was measured by ELISA (Figure 6C-F, right panel). In healthy controls, T_FH_ cells but not non-T_FH_ cells efficiently supported autologous memory B cells differentiated into total ASCs, as expected spike-specific ASCs and IgG were rarely observed (Figure 6C). T_FH_ cells from convalescents and vaccinees efficiently supported autologous memory B cells differentiated into both total ASCs and spike-specific ASCs and produced spike-specific IgG (Figure 6D-E). The third dose further enhanced the humoral immune responses, as more spike-specific ASCs and IgG were produced in comparison with before booster (Figure 6F). Moreover, the frequencies of spike-specific ASCs correlated with spike-specific IgG OD_450_ (Supporting Figure 11). Taken together, we conclude that T_FH_ cells, but not non-T_FH_ cells, were the major player in assisting antibody production and maintenance in SARS-CoV-2 infection and vaccination, which positively correlated with antibody responses (Figure 3D and Supporting Figure 7D-F; Figure 4F and Figure 5F).

**Figure 6.**
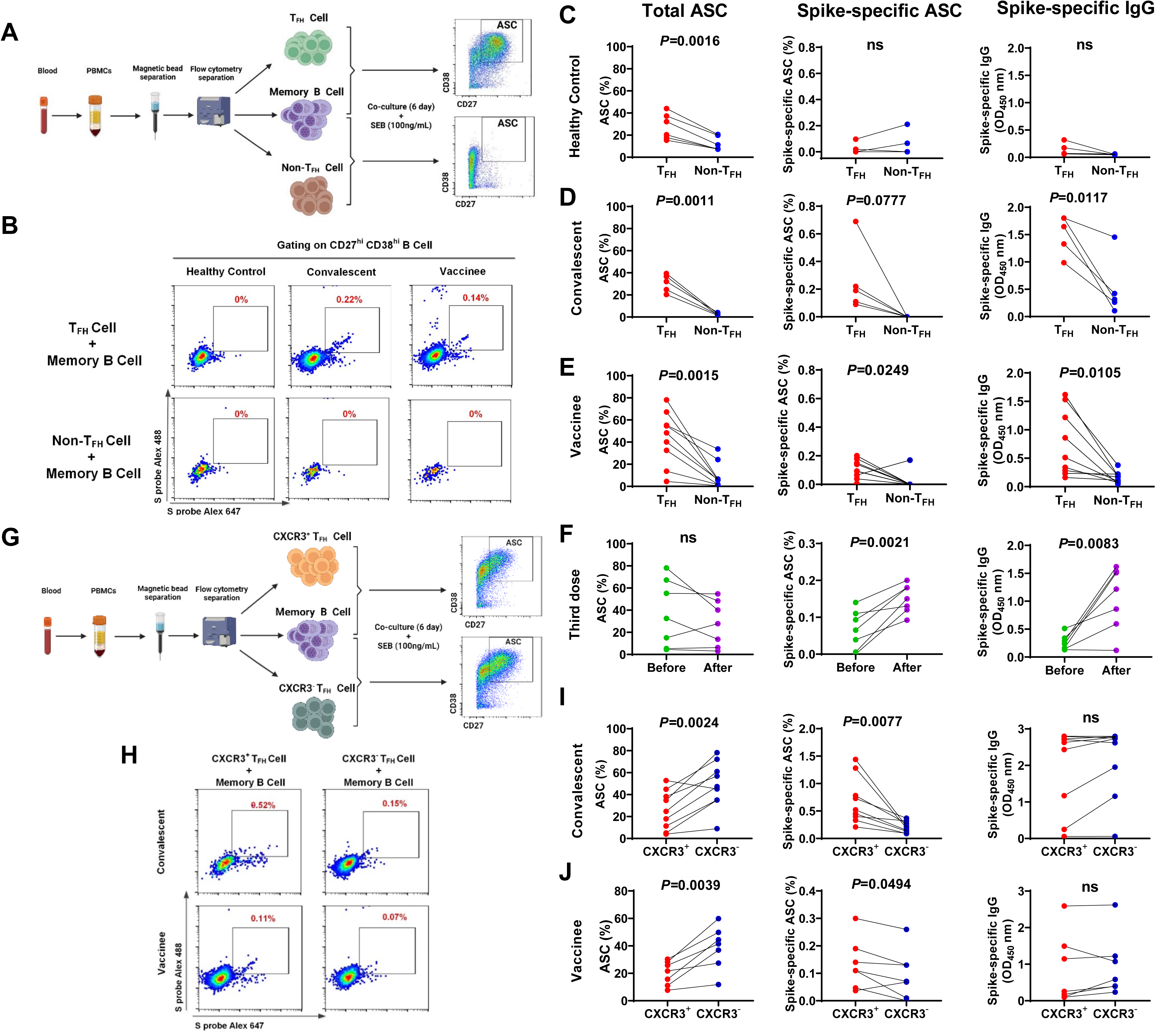
Spike-specific CXCR3^+^ T_FH_ cells exhibited superior capacity than spike-specific CXCR3^−^ T_FH_ cells in supporting ASC differentiation and antibody production. (A) Diagram of the coculture of T_FH_ cells or non-T_FH_ cells with autologous memory B cells. (B) Representative flow plots of spike-specific ASCs after coculture of T_FH_ cells or non-T_FH_ cells with memory B cells from healthy controls, convalescents, and vaccinees in the presence of SEB (100 ng/mL) for 6 days. (C-E) Comparison of total ASCs, spike-specific ASCs, and spike-specific IgG in supernatant between T_FH_ and non-T_FH_ cells after coculture with autologous memory B cells from healthy controls (n=6), convalescents (n=5), and vaccinees (n=9). (F) Total ASCs, spike-specific ASCs and spike-specific IgG before (n=7) and after (n=7) the third dose boost. (G) Diagram of coculture of CXCR3^+^ or CXCR3^−^ T_FH_ cells with autologous memory B cells. (H) Representative flow plots of spike-specific ASCs after coculture of CXCR3^+^ or CXCR3^−^ T_FH_ cells with memory B cells from convalescents and vaccinees in the presence of SEB (100 ng/mL) for 6 days. (I-J) Comparison of total ASCs, spike-specific ASCs, and spike-specific IgG in the supernatant between CXCR3^+^ T_FH_ cells and CXCR3^−^ T_FH_ cells after coculture with memory B cells from convalescents (n=9) and vaccinees (n=7) in the presence of SEB (100 ng/mL) for 6 days. In C-F and I-J, a paired t test was used to analyse the difference between two groups. *P*<0.05 was considered to be a two-tailed significant difference, ns means not significant.

According to the results above, we found that spike-specific CXCR3^+^ T_FH_ cells were associated with antibody magnitude and exhibited better responsiveness than spike-specific CXCR3^−^ T_FH_ cells in both natural infection and vaccination, but the functional difference of spike-specific CXCR3^+^ T_FH_ cells and spike-specific CXCR3^−^ T_FH_ cells was not clear. To further discriminate the functional role of spike-specific CXCR3^+^ T_FH_ cells and spike-specific CXCR3^−^ T_FH_ cells in supporting ASC differentiation and antibody production. Both bulk CXCR3^+^ and CXCR3^−^ T_FH_ cells were sorted from COVID-19 convalescents and vaccine recipients and cocultured with autologous memory B cells (Figure 6G). CXCR3^−^ T_FH_ cells were more efficient in supporting memory B-cell differentiation into total ASCs (Figure 6I and 6J, left panel); however, CXCR3^+^ T_FH_ cells showed superior capacity than CXCR3^−^ T_FH_ cells in supporting spike-specific ASC differentiation (Figure 6H; Figure 6I and 6J, middle panel), although we did not observe higher spike-specific IgG production in CXCR3^+^ T_FH_ cells than in CXCR3^−^ T_FH_ cells cocultured with autologous memory B cells, as expected (Figure 6I and 6J, right panel). This might be due to the lower proportions of spike-specific CXCR3^+^ T_FH_ cells within CXCR3^+^ T_FH_ cells than spike-specific CXCR3^−^ T_FH_ cells within CXCR3^−^ T_FH_ cells (Figure 2D and Figure 4E), although there was a higher responsiveness of spike-specific CXCR3^+^ T_FH_ cells than spike-specific CXCR3^−^ T_FH_ cells in convalescent and vaccine recipients (Figure 2D and Figure 4E). These findings further suggest that spike-specific T_FH_ cells, especially the spike-specific CXCR3^+^ subset, play a dominant role in assisting antibody production and maintenance in natural infection and vaccination.

## Discussion

In this study, we systemically investigated the longitudinal dynamics of spike-specific T_FH_ cell and antibody responses in individuals up to two years after COVID-19 recovery and those who received two and three doses of inactivated vaccine. We discovered that spike-specific T_FH_ cells, especially spike-specific CXCR3^+^ T_FH_ cells, played a dominant functional role in supporting high-quality antibody elicitation and maintenance in convalescents and inactivated vaccine recipients. Both natural infection and vaccination generated memory B cells, which could be recalled efficiently by spike-specific T_FH_ cells to differentiate into ASCs and produce spike-specific antibodies. These findings suggest that long-term humoral immunity could be generated and recalled, thus providing new insight into COVID-19 immunity and inspiring the evaluation of long-term protection.

Previous studies have shown that SARS-CoV-2 infection induces robust spike-specific T_H_1 and T_FH_ cell responses in the acute phase, which are maintained in the convalescent phase and persist for several months to a year(Dan *et al*., 2021; Marcotte et al., 2022; Rodda *et al*., 2021). Here, in the longitudinal analysis of circulating T_FH_ cells through a 2-year period in COVID-19 convalescents, we found that spike-specific T_FH_ cells rapidly declined from the 2^nd^ to the 5^th^ month and then remained stable for at least 12 months. The spike-specific CXCR3^+^ T_FH_ cell responses persisted for up to 24 months or more. Importantly, the decline in spike-specific PD-1^+^ active T_FH_ cells may account for the constraint memory T_FH_ cells after the 5^th^ month of recovery. In most symptomatic COVID-19 cases, robust GC reactions were demonstrated and found to be able to be maintained for at least six months(Laidlaw and Ellebedy, 2022; Poon *et al*., 2021). GC T_FH_ cells assist B-cell maturation and differentiation into high-affinity memory B cells and long-lived plasma cells. The long-term persistence of circulating T_FH_ cell responses may be attributed to active GC reactions in lymph nodes after recovery.

In line with the dynamics of T_FH_ cells, spike-specific IgG, IgG1, IgG3, and IgA antibody endpoint titers as well as neutralization titers showed a sharp decline from the 2^nd^ to the 5^th^ month and then were maintained in the majority of convalescents at the level for at least 24 months, longer than the 16 months reported recently(Yang *et al*., 2022). The sharp decrease was most likely due to the short half-life of serum antibodies and ASCs(Dan *et al*., 2021). ASCs normally decay within a few weeks, and only a population of long-lived plasma cells can live from several months to years(Hammarlund et al., 2017). IgA has been shown to play a dominant role in neutralization in early infection, although it is short-lived and declines rapidly after infection(Sterlin *et al*., 2021). Here, we found that spike-specific IgA antibodies were maintained for 24 months, although at a lower level, much longer than a previous observation that IgA was detectable six months after infection, which may also contribute to long-term protection(Roltgen and Boyd, 2021). Antibody maturation is a relatively slow process that is supported by T_FH_ cells in the GC(Crotty, 2019). Our results showed that spike-specific IgG antibody maturation peaked at the 5^th^-8^th^ month after infection, while spike-specific IgG3 and IgA became fully mature at the 12^th^ month. These data indicated that long-term T_FH_ cell responses are required in COVID-19 convalescents to gradually support high-quality antibody maturation and maintenance.

Inactivated vaccines have been widely used and proven to elicit robust and broad humoral immunity by the two-dose immunization procedure, which could protect against infection from VOC infection or severe disease(Chen et al., 2022; Liu et al., 2022). In this study, we also found that inactivated vaccine immunization mounted humoral immunity in a similar pattern to natural infection, but the magnitudes of antibody and T_FH_ cell responses were significantly lower. Although neutralizing antibody peaked at 14 days after the second dose and antibody matured at 3 months after the first dose, spike-specific T_FH_ cells could be observed at 14 days after the first dose, earlier than the appearance of neutralizing antibody. However, the neutralizing antibody significantly declined at 6 months after immunization. Based on the memory immunity generated by the two-dose injection, the third dose booster significantly magnified the neutralizing antibody titers, along with increased antibody maturation and responsive T_FH_ cells. These results were consistent with several recent studies on inactivated vaccine showing that a third-dose immunization increased the antibody neutralizing effect for SARS-CoV-2 and some VOCs, proving the enhancement and necessity of the third-dose booster(Liu *et al*., 2022; McMenamin et al., 2022). However, the T_FH_ cell and antibody responses elicited by inactivated vaccines seem to last a short period of time, different from natural infections, which elicit a lasting and relatively stable immune memory response(Marcotte *et al*., 2022). In addition, current intramuscular injection immunization only mounted a poor IgA response, which is important for mucosal immunity in the respiratory tract(Liu *et al*., 2022). Together, to maintain recallable immunity, heterogonous sequential immunization may be a strategy for improving immunization efficacy, especially intranasal immunization to elicit local mucosal immunity.

The long-lived antibody response is mainly supported by T_FH_ cells in natural infection and vaccination(Laidlaw and Ellebedy, 2022; Roltgen et al., 2022). We found that SARS-CoV-2 infection and vaccination induced robust spike-specific CD4^+^ T cells, comprising T_H_1 and T_FH_ cell responses, consistent with other studies, thus laying the foundation for long-term immunity(Chen *et al*., 2022; Liu *et al*., 2022). Longitudinal analysis showed that spike-specific T_FH_ and non-T_FH_ cell responses were maintained for at least 12 months and were more stable in COVID-19 convalescent than inactivated vaccine recipients, although the dynamics were similar in both cohorts. Notably, spike-specific T_FH_ cells, especially spike-specific CXCR3^+^ T_FH_ cells, positively correlated with antibody responses at the 5^th^ month in convalescents, in which the quantity and quality of antibodies were well balanced. Although spike-specific T_FH_ cells did not correlate with the neutralizing antibody titer in the vaccine recipients, the ratio of spike-specific CXCR3^+^/CXCR3^−^ T_FH_ cells at Day 14 after the first dose, representative of T_H_1-like T_FH_ cell bias, positively correlated with the neutralizing titer on Day 45 (Day 14 after the second dose). In addition, the T_FH_ cells, not non-T_FH_ cells, elicited by the two-dose immunization were associated with the antibody magnitude after the third dose. Functionally, T_FH_ cells could recall autologous memory B cells to differentiate into total ASCs, spike-specific ASCs and antibodies, but non-T_FH_ cells could not stimulate a detectable level of ASCs. Therefore, circulating spike-specific T_FH_ cells are a surrogate of bona fide GC T_FH_ cells that support spike-specific ASC differentiation and antibody production in natural infection and vaccination(Morita *et al*., 2011). Of note, our study did not exclude the possibility that non-T_FH_ cells supported short-lived plasmablast differentiation and produced low affinity antibodies in the very early acute phase to constrain infection rapidly, as large amounts of antibodies could be produced by extrafollicular B cells in some severe cases(Woodruff et al., 2020).

SARS-CoV-2 infection-induced circulating T_FH_ cells exhibited a clear phenotypic bias towards a CCR6^+^ CXCR3^−^ phenotype, and these subsets comprised the large proportions of spike-specific T_FH_ cells in natural infection(Dan *et al*., 2021; Juno *et al*., 2020; Rodda *et al*., 2021; Rydyznski Moderbacher *et al*., 2020). We and others have previously shown that the CXCR3^+^ T_FH_ cell subset is increased in convalescents and positively correlated with the spike-specific antibody response(Gong *et al*., 2020; Juno *et al*., 2020; Zhang *et al*., 2021). The association of T_H_1-like (CXCR3^+^) T_FH_ cells with the quantity and quality of antibody was also characterized in influenza vaccine recipients and in other chronic viral infections(Bentebibel *et al*., 2016; Bentebibel *et al*., 2013; Koutsakos et al., 2018; Niessl *et al*., 2020; Zhang *et al*., 2019). Furthermore, we have demonstrated that CXCR3^+^ and CXCR3^−^ T_FH_ subsets have distinct phenotypes and functions in HCV infection(Zhang *et al*., 2019). Here, spike-specific CXCR3^+^ T_FH_ cells showed an active status and maintained responsiveness for a longer time than spike-specific CXCR3^−^ T_FH_ cells in convalescents. However, the functional significance of CXCR3^+^ T_FH_ cells in SARS-CoV-2 infection and vaccination is extremely unknown. Coculture of CXCR3^+^ T_FH_ or CXCR3^−^ T_FH_ cells with autologous memory B cells from COVID-19 convalescents and vaccinees showed that CXCR3^+^ T_FH_ cells exhibited superior capacity than CXCR3^−^ T_FH_ cells in supporting spike-specific memory B cells differentiated into ASCs, confirming that spike-specific CXCR3^+^ T_FH_ cells played a dominant function in assisting the antibody response in natural infection and vaccination, although the proportions of spike-specific CXCR3^+^ T_FH_ cells were lower than those of CXCR3^−^ T_FH_ cells.

The T_H_1-polarizing conditions of a viral infection usually result in the predominant generation of T_H_1-like T_FH_ cells, such as influenza vaccination, following live-attenuated yellow fever vaccination and HCV and Zika virus infection(Bentebibel *et al*., 2016; Bentebibel *et al*., 2013; Liang et al., 2019; Martin-Gayo *et al*., 2017; Zhang *et al*., 2019). Given the importance of T_H_1-like T_FH_ cells in supporting the production of high-quality antibodies and the longevity of secreting cells, strategies to promote T_H_1-like T_FH_ cell polarization would benefit SARS-CoV-2 vaccine improvement. In this study, we found that spike-specific CXCR3^+^ T_FH_ cells show long-term persistence and play a dominant role in supporting the antibody response. These findings will inform vaccine design towards long-term protection against SARS-CoV-2 infection.

## Materials and Methods

### Study subjects

This study included two cohorts, convalescents and vaccinated subjects. Thirty-seven COVID-19 convalescent subjects were from The Central Hospital of Shaoyang, Hunan Province, China, and all were followed up to 24 months after symptom onset (Supporting Table 1). Another 37 vaccinated subjects were from The First People’s Hospital of Chenzhou, Hunan Province, China. All received two doses of inactivated vaccine (Sinovac, China), and 24 received the third-dose vaccination (Supporting Table 2). Each participant signed a written consent form. The study protocol was approved by the Institutional Ethical Review Board of The Central Hospital of Shaoyang (V.1.0, 20200301) and the First People’s Hospital of Chenzhou (V.3.0, 2021001). Blood samples of COVID-19 convalescents were collected at the 2^nd^, 5^th^, 8^th^, 12^th^ and 24^th^ months after COVID-19 symptom onset. Blood samples of vaccine recipients were collected before vaccination and 14, 28, 45, 90, and 180 days after inoculation with the first dose; blood samples from 24 recipients were taken before and 14 days after the third dose vaccination (Supporting Table 1 & Table 2). Seventeen healthy individuals before the COVID-19 pandemic were used as the negative control. PBMCs and plasma were isolated and stored in liquid nitrogen in a −80 °C freezer.

### Endpoint titer of antibody

The endpoint titers of spike-specific antibodies were determined by measuring the binding activity of serially diluted plasma to the SARS-CoV-2 spike protein using ELISA. In brief, 96-well plates (Corning, NY, USA) were coated with SARS-CoV-2 spike protein (SARS-CoV-2 S1+S2_ECD, 200 ng/well) (Sino Biological, Beijing, China) in PBS and incubated at 4 °C overnight. The plates were washed five times with PBS-T (0.05% Tween-20 in PBS) and then blocked with blocking buffer (2% FBS and 2% BSA in PBS-T) for 30 min. Two-fold serial dilutions of plasma, starting from a 1:20 dilution, were added to the 96-well plates in triplicate (100 µl/well) and incubated for 1 h at room temperature. Spike-specific antibodies were detected using horseradish peroxidase (HRP)-conjugated anti-human IgG (Jackson ImmunoResearch, PA, USA), IgG1, IgG3 (BaiaoTong Experiment Center, Luoyang, China), and IgA (Thermo Fisher Scientific, Waltham, MA, USA). Plasma from healthy subjects was collected before the COVID-19 pandemic as a negative control, and SARS-CoV-2 spike RBD-specific monoclonal antibody was generated in the laboratory and used as a positive control. Optical density at 450 nm (OD_450_) was measured for each reaction, and an OD_450_ of three-fold above the cut-off value was considered a positive readout. The highest dilution showing a positive readout was defined as the endpoint titer of the antibody, and the data were logarithmically transformed.

### Antibody avidity assay

The avidity of spike-specific antibodies (IgG, IgG1, IgG3 and IgA) was measured using a modified 2-step approach that we described previously(Zhang *et al*., 2021). In the first step, plasma dilutions were optimized to obtain an OD_450_ value within the range of 0.5-1.5 to ensure a linear measurement of the antibody avidity. The second step was an ELISA but included an elution procedure of 1 M NaSCN. These measurements were performed in triplicate. The avidity index of an antibody was calculated as OD_NaSCN 1M_/OD_NaSCN 0M_ × 100%.

### Antibody neutralization assay

The neutralization activity of plasma was determined by the reduction in luciferase expression after pseudotyped virus infection of Huh7 cells, as described previously(Zhang *et al*., 2021). In brief, SARS-CoV-2 pseudotyped virus was incubated in duplicate with serial dilutions of plasma samples (six dilutions: 1:30; 1:90; 1:270; 1:810; 1:2430; 1:7290) at 37 °C for 1h. Then, freshly trypsinized cells were added at 5% CO_2_ and incubated at 37 °C for 24 h, and the luminescence was measured. In parallel, control wells with virus only or cells only were included in six replicates. The background relative light unit (RLU) (wells with cells only) was subtracted from each determination. Plasma from healthy controls was used as a negative control. Plasma from guinea pigs immunized with the SARS-CoV-2 spike protein was used as a positive control. The 50% inhibitory dilution (ID_50_) was defined as the plasma dilution, of which the RLU was reduced by 50% compared with the control solution wells (virus + cells). The cut-off value was defined as ID_50_=30, and ID_50_>30 was considered to have a neutralization effect.

### Antigen-specific T_FH_ cell assay

To analyse spike-specific T_FH_ cells, a CD154 (CD40 L) assay was used to assess the response of circulating T_FH_ cells upon stimulation. In brief, cryopreserved PBMCs were thawed and allowed to recover in complete RPMI 1640 medium in 5% CO_2_ at 37 °C overnight. PBMCs (1×10^6^) were stimulated with SARS-CoV-2 spike protein (S1+S2_ECD, 5 μg/ml, Sino Biological, Beijing, China) or BSA (5 μg/ml, SigmaLJAldrich, St. Louis, MO, USA) in 5% CO_2_ at 37 °C for 24 h, PE mouse anti-human CD154 (24-31) (BioLegend, San Diego, CA, USA) was added during the stimulation. Concanavalin A (Con A, 5 μg/ml, SigmaLJAldrich, St. Louis, MO, USA) was used as a positive control. In parallel, PBMCs from healthy controls were stimulated under the same conditions. After stimulation, the cells were labelled with a LIVE/DEAD^®^ Fixable Blue Dead Cell Stain Kit (Thermo Fisher Scientific, Waltham, MA, USA) to distinguish dead cells and then treated with Fc Block (BioLegend, San Diego, CA, USA) to block nonspecific binding. The treated PBMCs were stained with antibodies that had been pretitrated to an optimized dilution and fluorescently labelled in 96-well V-bottom plates at 4 °C for 30 min. The fluorescent labelled antibodies were BUV737 mouse anti-human CD4 (SK3), PE mouse anti-human CXCR3 (1C6) (BD Biosciences, Franklin Lake, NJ, USA), FITC mouse anti-human PD-1 (EH12.2H7) (BioLegend, San Diego, CA, USA), and PE-eFluor 610 mouse anti-human CXCR5 (MU5UBEE) (Thermo Fisher Scientific, Waltham, MA, USA). Samples were loaded onto a MoFlo XDP Flow Cytometer (Beckman Coulter, Brea, CA, USA) immediately after antibody staining. The cells were gated with the strategy (Supporting Figure 1) as spike-specific T_FH_ cells (CD154^+^ CXCR5^+^ CD4^+^ T cells), spike-specific CXCR3^+^ T_FH_ cells (CD154^+^ CXCR3^+^ CXCR5^+^ CD4^+^ T cells), spike-specific CXCR3^−^ T_FH_ cells (CD154^+^ CXCR3^−^ CXCR5^+^ CD4^+^ T cells), active T_FH_ cells (CD154^+^ PD-1^+^ CXCR5^+^ CD4^+^ T cells), and spike-specific non-T_FH_ cells (CD154^+^ CXCR5^−^ CD4^+^ T cells). The responsiveness capacity presented as the stimulation index (SI: spike-responsive versus BSA-responsive [baseline]). The gating of cell populations was based on the mean fluorescence intensity “minus one” (FMO) and unstained control. All data were analysed using FlowJo 10.0 software (Tree Star, San Carlos, CA, USA).

### T_FH_ and memory B-cell coculture

To test the functional role of T_FH_ cells, CXCR3^+^ and CXCR3^−^ T_FH_ cell subsets, and non-T_FH_ cells in supporting ASC differentiation and antibody production, these cells were cocultured with autologous memory B cells. In detail, CD4^+^ T cells and CD19^+^ B cells were primarily sorted from PBMCs of convalescents and vaccinees by CD4 and CD19 MicroBeads (Miltenyi Biotec, Bergisch Gladbach, Germany), respectively. Purified CD4^+^ T cells and CD19^+^ B cells were sorted into T_FH_ cells (CXCR5^+^ CD4^+^ T cells), non-T_FH_ cells (CXCR5^−^ CD4^+^ T cells), CXCR3^+^ T_FH_ cells (CXCR3^+^ CXCR5^+^ CD4^+^ T cells), CXCR3^−^ T_FH_ cells (CXCR3^−^ CXCR5^+^ CD4^+^ T cells), and memory B cells (CD20^+^ CD27^+^ B cells) by FACS. CD4^+^ T cells were labelled with a LIVE/DEAD^®^ Fixable Blue Dead Cell Stain Kit (Thermo Fisher Scientific, Waltham, MA, USA) to distinguish dead cells in a 15 ml centrifuge tube at 4 °C for 30 min and then stained with BUV737 mouse anti-human CD4 (SK3), PE mouse anti-human CXCR3 (1C6) (BD Biosciences, Franklin Lake, NJ, USA), and PE-eFluor 610 mouse anti-human CXCR5 (MU5UBEE) (Thermo Fisher Scientific, Waltham, MA, USA). CD19^+^ B cells were stained with mouse anti-human CD20 PE (2H7), mouse anti-human PE-Cy™7 CD27 (M-T271) (BioLegend, San Diego, CA, USA). Sorted T_FH_, non-T_FH_, CXCR3^+^ T_FH_, and CXCR3^−^ T_FH_ cells (5×10^4^ cells for each type) were cocultured with autologous memory B cells (5×10^4^ cells) in the presence of 100 ng/ml *staphylococcal enterotoxin* B (SEB) (Toxin Technology, Sarasota, FL, USA) and RPMI 1640 medium supplemented with 10% FBS in 96-well U-bottom plates for 6 days. After coculture, spike-specific IgG in the supernatant was determined by ELISA. Antibody-secreting cells (ASCs) (CD4^−^ CD27^hi^ CD38^hi^ cells) and spike-specific ASCs (spike-probe FITC^+^ and spike-probe ALEX647^+^ CD4^−^ CD27^hi^ CD38^hi^ cells) were analysed by flow cytometry. Spike-specific ASCs were intracellularly stained. All data analyses were performed with FlowJo 10.0 software (Tree Star, San Carlos, CA, USA).

## Statistical analysis

All data were analysed by the Kolmogorov–Smirnov test for normal distribution. When variables were nonnormally distributed, data were expressed as the median ± IQR (interquartile range), and Mann–Whitney U tests were used to analyse two independent variables. For paired sample comparison, data were expressed as each individual value or mean ± SEM (standard error of mean), and paired t tests were used to analyse the differences between two groups. Differences among multiple groups were compared with one-way analysis of variance, and Tukey’s test was used between two groups at the same time point. Spearman’s rank correlation coefficient was used to measure the correlation between two different variables. Analyses of the data were performed using SPSS v.26 and GraphPad Prism v.8.0. All numerical data shown in this study were collected from at least three independent experiments.

## Acknowledgements

We thank all of the participants. This work was supported by the National Natural Science Foundation of China (82061138020, 82102365), Natural Science Foundation of Hunan Province of China (2021JJ40006, 2022JJ30095), Educational Commission of Hunan Province of China (21A0529), Key Research and Development Project of Chenzhou City, Hunan Province (ZDYF2020010, ZDYF2020007, zdyf201920, zdyf201921), Key Project of The First People’s Hospital of Chenzhou (N2019-002) and SC1-PHE-CORONAVIRUS-2020: “Advancing knowledge for the clinical and public health response to the 2019-nCoV epidemic” from the European Commission (CORONADX, no. 101003562) (Y.-P.L).

## Author contributions

X.Q., W. L, Y.-P.L. and Y.W. contributed to the study design and data interpretation. Q.W., Q.J.W., T.X. and T.B. contributed to patient recruitment and sample collection and processing. Z.L., Y.H., J.C. and J.Y. contributed to the serum antibody binding and avidity experiments. F.C., Y.L. and S.T. contributed to pseudotyped virus production and antibody neutralization experiments. J. Z, X.Z., B.L. and B.W. contributed to all T_FH_ experiments and data analysis. X.Q., J.Z., R.H., B.L. and X.Z. drafted the manuscript. X.Q., W.L. and Y.-P.L., V.T. and Y.W. contributed to critical revision of the manuscript for important intellectual content. X.Q., W.L. and Y.-P.L. provided supervision. All authors met authorship criteria and approved the publication.

## Supporting Figure Legends

**Supporting Figure 1. Gating strategy of spike-specific T_FH_ and non-T_FH_ cells**

Gating strategy of spike-specific T_FH_ cells (CD154^+^ CXCR5^+^ CD4^+^ T cells), spike-specific CXCR3^+^ T_FH_ cells (CD154^+^ CXCR3^+^ CXCR5^+^ CD4^+^ T cells), spike-specific CXCR3^−^ T_FH_ cells (CD154^+^ CXCR3^−^ CXCR5^+^ CD4^+^ T cells), active T_FH_ cells (CD154^+^ PD-1^+^ CXCR5^+^ CD4^+^ T cells), and non-T_FH_ cells (CD154^+^ CXCR5^−^ CD4^+^ T cells).

**Supporting Figure 2. Longitudinal analysis of non-T_FH_ cell responses in COVID-19 convalescents**

(A) The frequencies of spike-specific non-T_FH_ cells at the 2^nd^ month (n=20), 5^th^ month (n=18), 8^th^ month (n=20), 12^th^ month (n=20), and 24^th^ month (n=5) after COVID-19 illness onset, as well as healthy controls (n=17) upon BSA or spike protein stimulation. A paired t test was used to analyse the difference in BSA and spike protein stimulation. (B) Kinetics of spike-specific non-T_FH_ cell responses upon BSA or spike protein stimulation and SI at the indicated time points. One-way analysis of variance was used to compare the differences among multiple groups, and Tukey’s multiple-comparisons test was used to compare differences within groups. The data are presented as the median ± IQR (25%– 75%). *P*<0.05 was considered to be a two-tailed significant difference, ns means not significant.

**Supporting Figure 3. Differential responsiveness of spike-specific PD-1+ CXCR3+ and PD-1+ CXCR3− TFH cells in COVID-19 convalescents**

Frequencies of spike-specific PD-1^+^ CXCR3^+^ (A) and PD-1^+^ CXCR3^−^ T_FH_ cells (B) at the 2^nd^ month (n=20), 5^th^ month (n=18), 8^th^ month (n=20), 12^th^ month (n=20) and 24^th^ month (n=5) as well as healthy controls (n=17) upon BSA or spike protein stimulation. (C) Dynamics of spike-specific PD-1^+^ CXCR3^+^ and PD-1^+^ CXCR3^−^ T_FH_ cell responses at the indicated time points, as well as the SIs of spike-specific PD-1^+^ CXCR3^+^ and PD-1^+^ CXCR3^−^ T_FH_ cells. Data are the mean ± SEM. In A-C, a paired t test was used to analyse the difference between two groups. * *P*<0.05; ** *P*<0.01; *** *P*<0.001; *** *P*<0.0001. *P*< 0.05 was considered to be a two-tailed significant difference, ns means not significant.

**Supporting Figure 4. Longitudinal analysis of non-T_FH_ cell responses in COVID-19 convalescents**

Frequencies of spike-specific CXCR3^+^ (A) and CXCR3^−^ non-T_FH_ cells (B) at the 2^nd^ month (n=20), 5^th^ month (n=18), 8^th^ month (n=20), 12^th^ month (n=20), and 24^th^ month (n=5) as well as healthy controls (n=17) upon BSA or spike protein stimulation. A paired t test was used to analyse the difference in BSA and spike protein stimulation. (C) Dynamics of spike-specific CXCR3^+^ and CXCR3^−^ non-T_FH_ cells at the indicated time points upon BSA or spike protein stimulation as well as their SIs. One-way analysis of variance was used to compare the differences among multiple groups, and Tukey’s multiple-comparisons test was used to compare differences within the groups. Data are presented as the median ± IQR (25%–75%). (D) Comparison of spike-specific CXCR3^+^ and CXCR3^−^ non-T_FH_ cell responses as well as SIs at the indicated time points. Data are the mean ± SEM. A paired t test was used to analyse the difference between spike-specific CXCR3^+^ and CXCR3^−^ non-T_FH_ cells. * *P*<0.05; ** *P*<0.01; *** *P*<0.001; **** *P*<0.0001. *P*< 0.05 was considered to be a two-tailed significant difference, ns means not significant.

**Supporting Figure 5. Correlations of spike-specific T_FH_ cells and non-T_FH_ cells in COVID-19 convalescents**

Correlations of spike-specific T_FH_ cells and spike-specific non-T_FH_ cells (A), spike-specific CXCR3^+^ T_FH_ cells and spike-specific CXCR3^+^ non-T_FH_ cells (B), and spike-specific CXCR3^−^ T_FH_ cells and spike-specific CXCR3^−^ non-T_FH_ cells (C) in COVID-19 convalescents at the 2^nd^ month (n=20), 5^th^ month (n=18), 8^th^ month (n=20), 12^th^ month (n=20), and 24^th^ month (n=5). In A, B and C, Spearman’s rank correlation coefficient was used to describe the association between spike-specific T_FH_ cells and spike-specific non-T_FH_ cells. *P*< 0.05 was considered to be a two-tailed significant difference.

**Supporting Figure 6. Correlations of spike-specific antibody endpoint titers and neutralization titers in COVID-19 convalescents**

Correlation of spike-specific IgG (A), IgG1 (B), IgG3 (C), and IgA (D) endpoint titers and neutralization in COVID-19 convalescents at the 2^nd^ month (n=25), 5^th^ month (n=25), 8^th^ month (n=25), 12^th^ month (n=24), and 24^th^ month (n=5). Spearman’s rank correlation coefficient was used to describe the association between the spike-specific endpoint titer and neutralization titers. *P*< 0.05 was considered to be a two-tailed significant difference.

**Supporting Figure 7. Correlations of spike-specific IgG1 endpoint titers and spike-specific T_FH_ cells or spike-specific non-T_FH_ cells in COVID-19 convalescents**

(A-C) Correlation analysis of spike-specific IgG1 endpoint titers with spike-specific T_FH_ cells (A), spike-specific CXCR3^+^ T_FH_ cells (B), and spike-specific CXCR3^−^ T_FH_ cells (C) in COVID-19 convalescents at the 5^th^ month (n=18). (D-F) Correlation analysis of IgG1 endpoint titers with spike-specific non-T_FH_ cells (D), spike-specific CXCR3^+^ non-T_FH_ cells (E), and spike-specific CXCR3^−^ non-T_FH_ cells (F) in COVID-19 convalescents at the 5^th^ month (n=18). Spearman’s rank correlation coefficient was used to describe the association between IgG1 endpoint titers and T_FH_ cell subsets or non-T_FH_ cell subsets. *P*< 0.05 was considered to be a two-tailed significant difference.

**Supporting Figure 8. Spike-specific PD-1^+^ T_FH_ cell subset responses in inactivated vaccine recipients**

(A-B) Spike-specific PD-1^+^ CXCR3^+^ T_FH_ cells (A) and PD-1^+^ CXCR3^−^ T_FH_ cell (B) responses upon spike stimulation at the indicated time points in vaccinees. (C) Comparison of spike-specific PD-1^+^ CXCR3^+^ T_FH_ cell and PD-1^+^ CXCR3^−^ T_FH_ cell responses as well as SI at the indicated time points. In A-C, a paired t test was used to analyse the differences between the two groups at the indicated times, and the data are the mean ± SEM. * *P*<0.05; ** *P*<0.01; *** *P*<0.001; **** *P*<0.0001. *P*< 0.05 was considered to be a two-tailed significant difference, ns means not significant.

**Supporting Figure 9. Spike-specific non-T_FH_ cell responses in inactivated vaccine recipients**

Spike-specific CXCR3^+^ non-T_FH_ cell (A) and CXCR3^−^ non-T_FH_ cell responses (B) upon spike stimulation at the indicated time points. (C) Comparison of spike-specific CXCR3^+^ non-T_FH_ cell and CXCR3^−^ non-T_FH_ cell frequencies as well as SIs. In A-C, a paired t test was used to analyse the difference between two groups at the indicated times, and the data are the mean ± SEM. * *P*<0.05; ** *P*<0.01; *** *P*<0.001; **** *P*<0.0001. *P*< 0.05 was considered to be a two-tailed significant difference, ns means not significant.

**Supporting Figure 10. A third dose boost promoted spike-specific non-T_FH_ cell responses in inactivated vaccine recipients**

(A) Frequencies of spike-specific non-T_FH_ cells, spike-specific CXCR3^+^ non-T_FH_ cells, and spike-specific CXCR3^−^ non-T_FH_ cell responses before and 14 days after the third dose boost upon BSA or spike protein stimulation (n=24). (B) SIs of spike-specific non-T_FH_ cells, spike-CXCR3^+^ non-T_FH_ cells, and spike-specific CXCR3^−^ non-T_FH_ cells before and 14 days after the third dose boost (n=24). (C) Comparison of SI between spike-specific CXCR3^+^ non-T_FH_ cells and spike-specific CXCR3^−^ non-T_FH_ cells before and 14 days after the third dose boost (n=24). In A-C, a paired t test was used to analyse the difference between two groups. *P*< 0.05 was considered to be a two-tailed significant difference, ns means not significant.

**Supporting Figure 11. Correlation of spike-specific ASCs with spike-specific IgG in supernatants after coculture**

Correlation of spike-specific ASCs and the spike-specific IgG OD_450_ after coculture of T_FH_ cells with autologous memory B cells from convalescents (n=5) and vaccinated subjects (n=14) in the presence of SEB (100 ng/mL) for 6 days. Spearman’s rank correlation coefficient was used to measure the correlation between two different variables. *P*< 0.05 was considered to be a two-tailed significant difference.

