## Supporting Figures for "Spike-specific CXCR3^+^ T_FH_ cells play a dominant functional role in supporting antibody responses in SARS-CoV-2 infection and vaccination"

### Supporting Figure 1

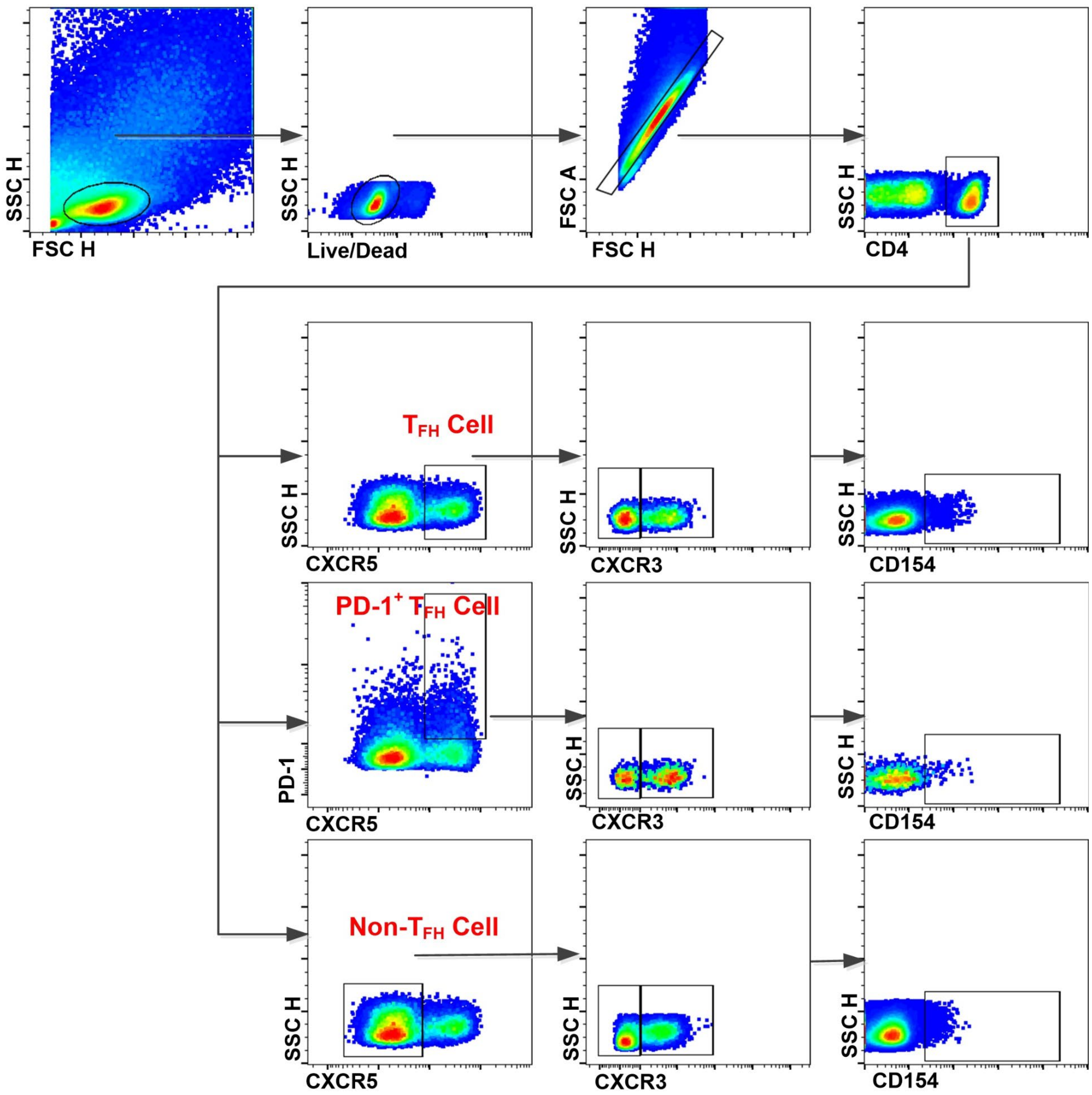

**Supporting Figure 2**

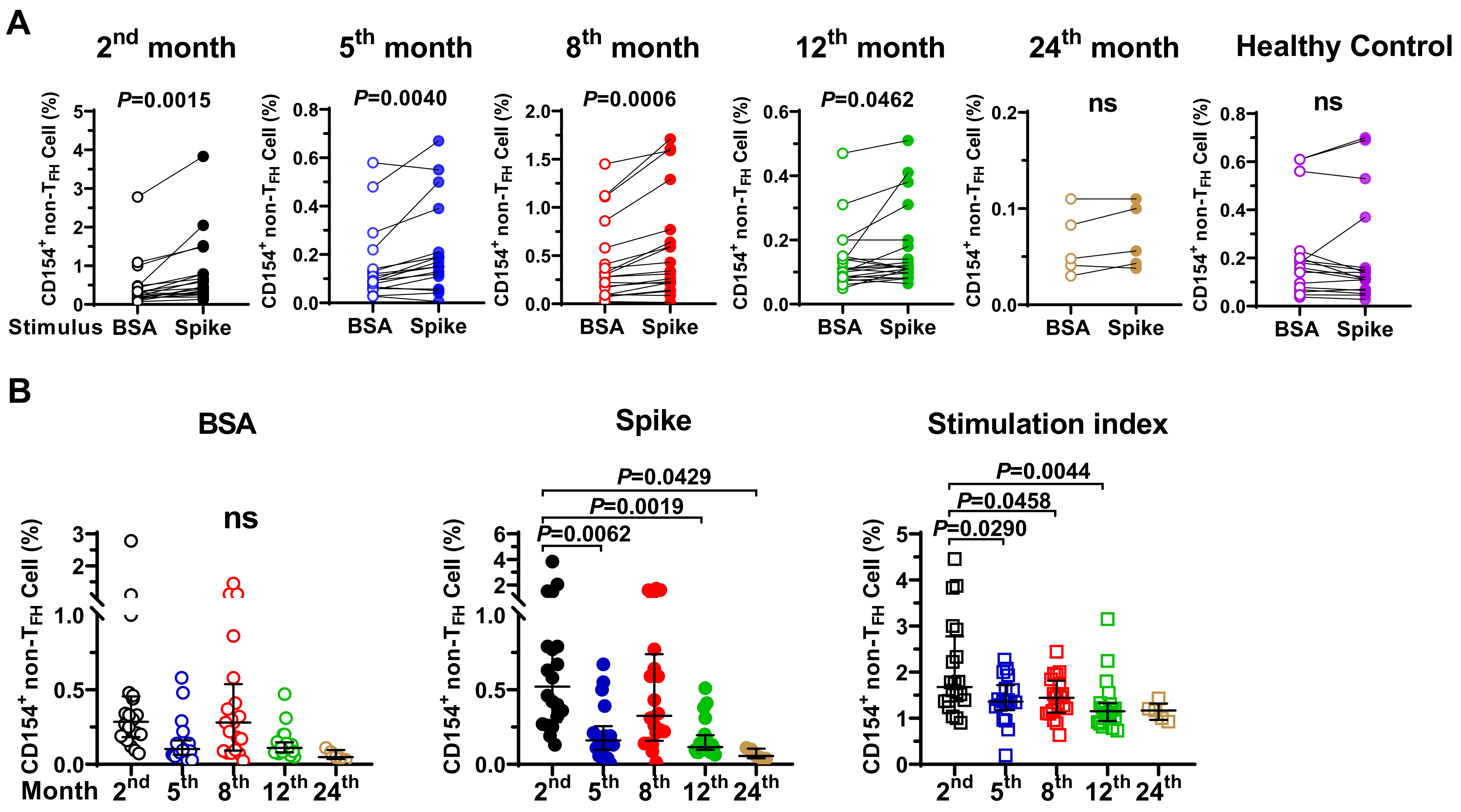

**Supporting Figure 3**

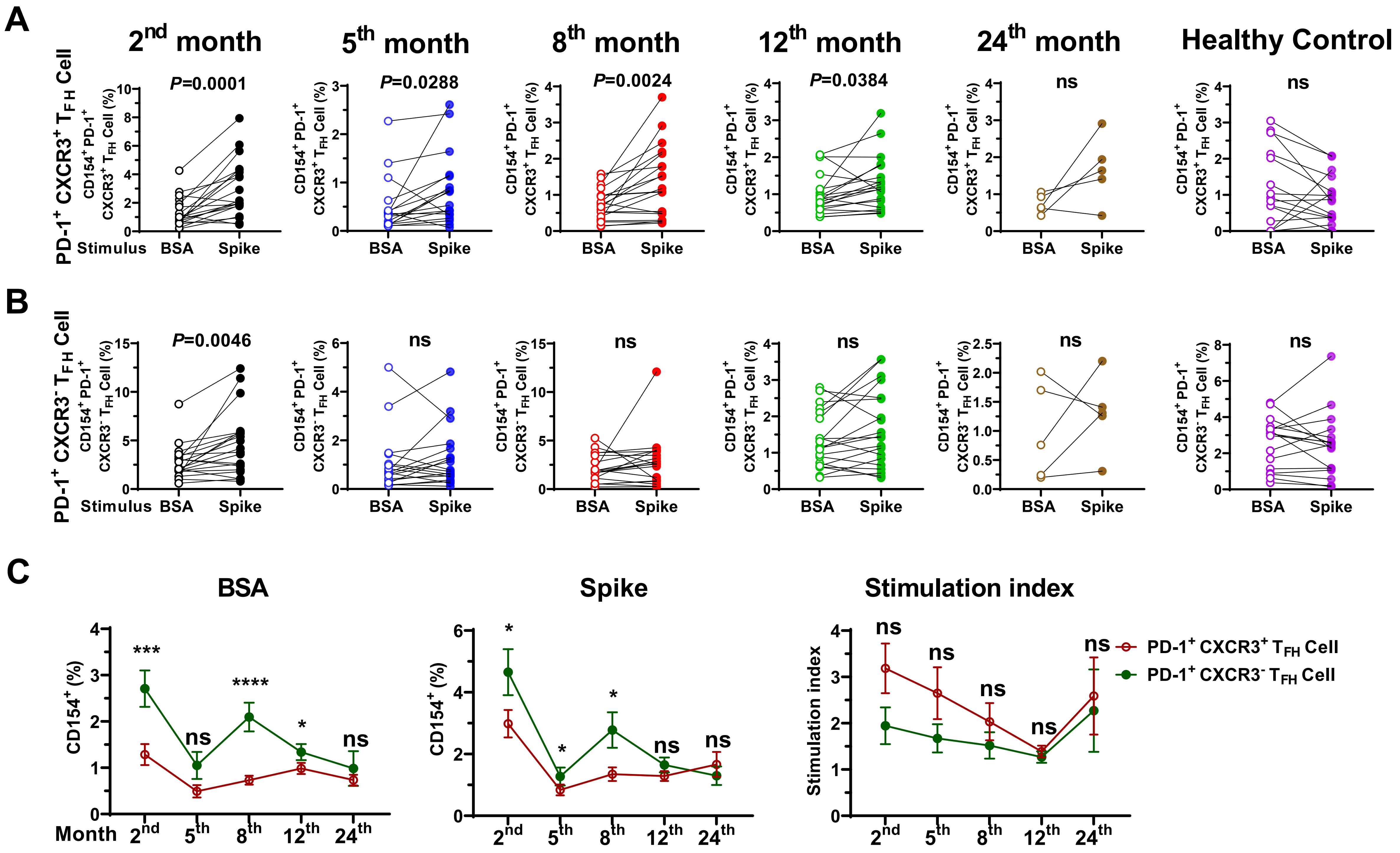

### Supporting Figure 4

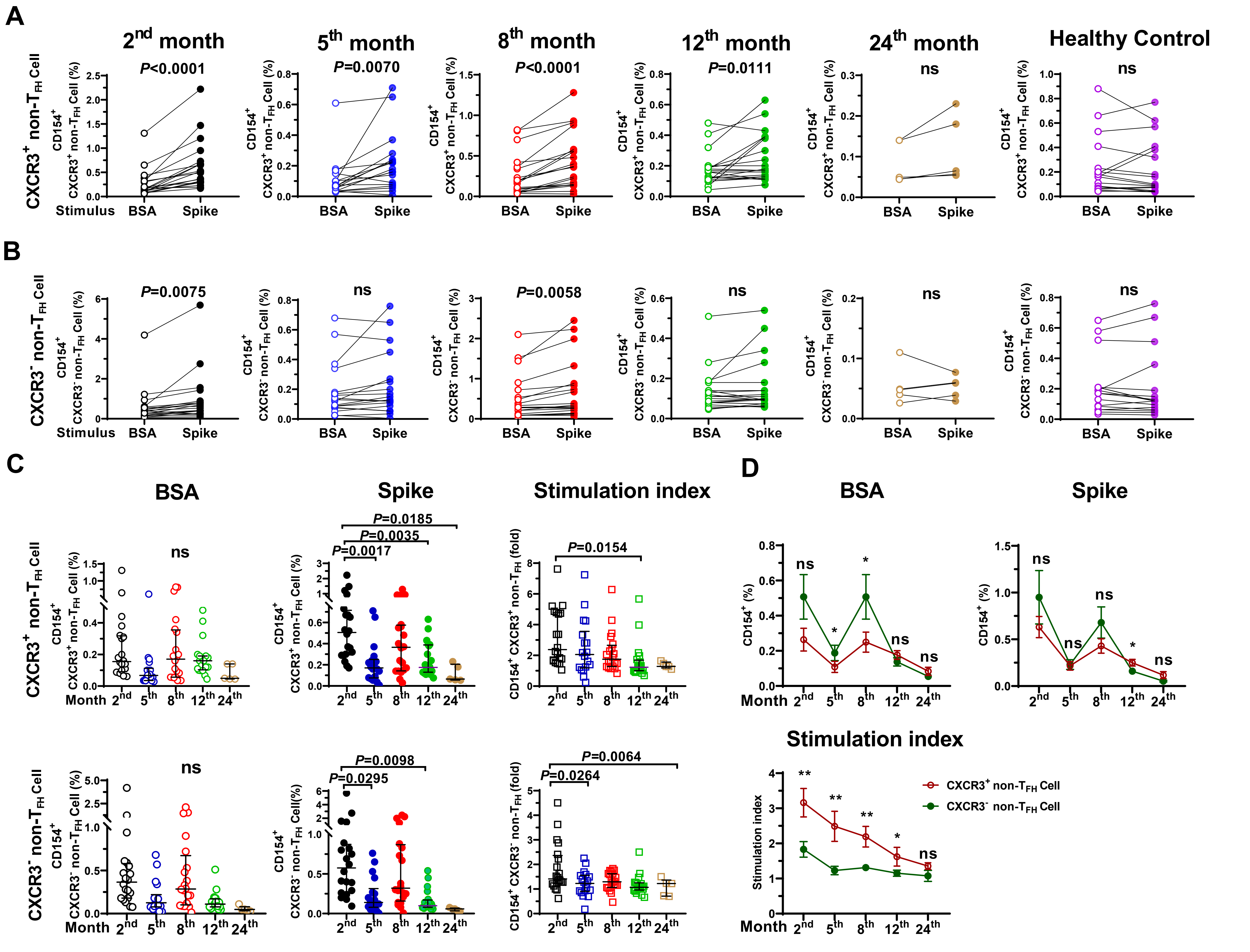

### Supporting Figure 5

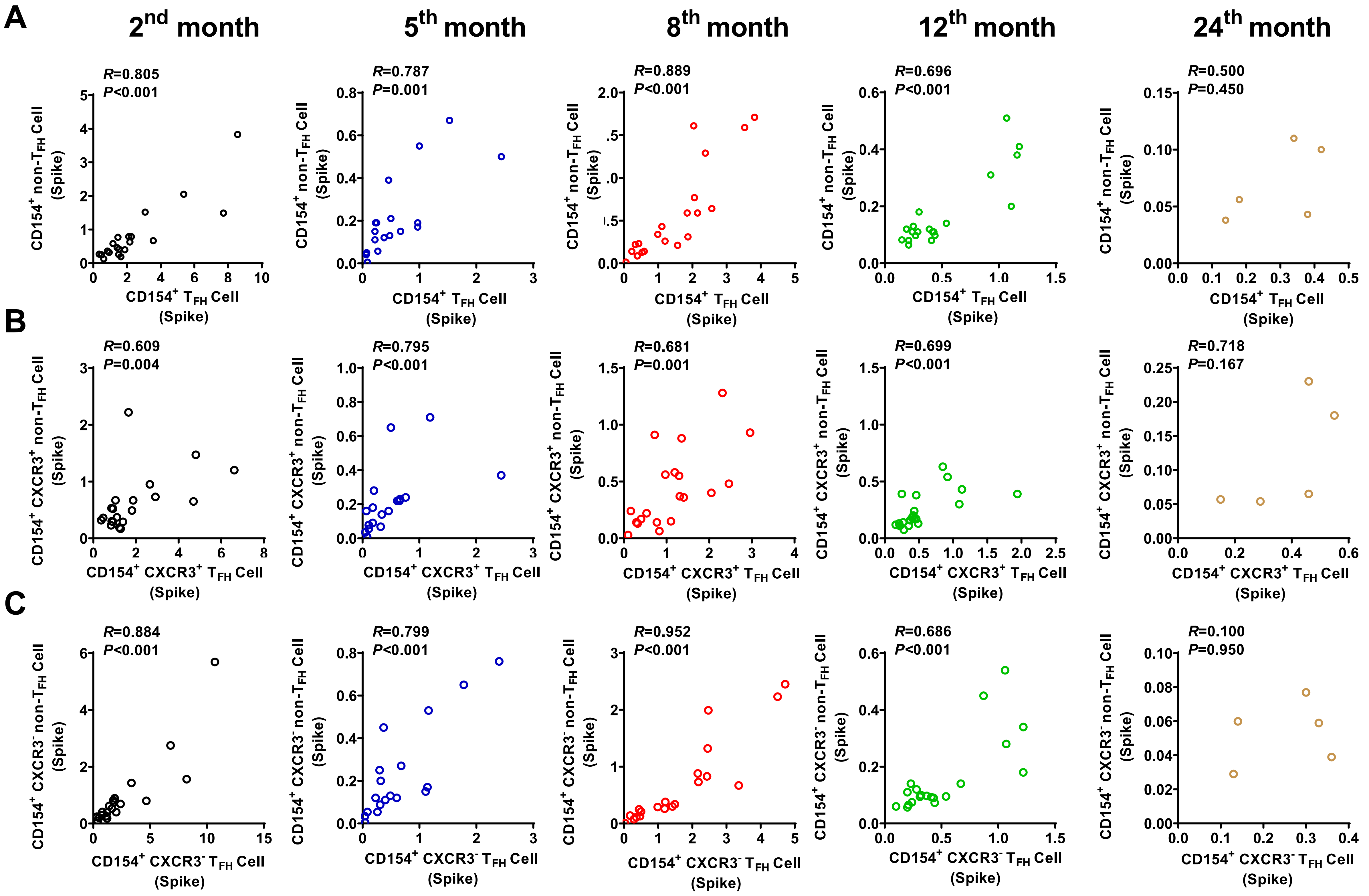

### Supporting Figure 6

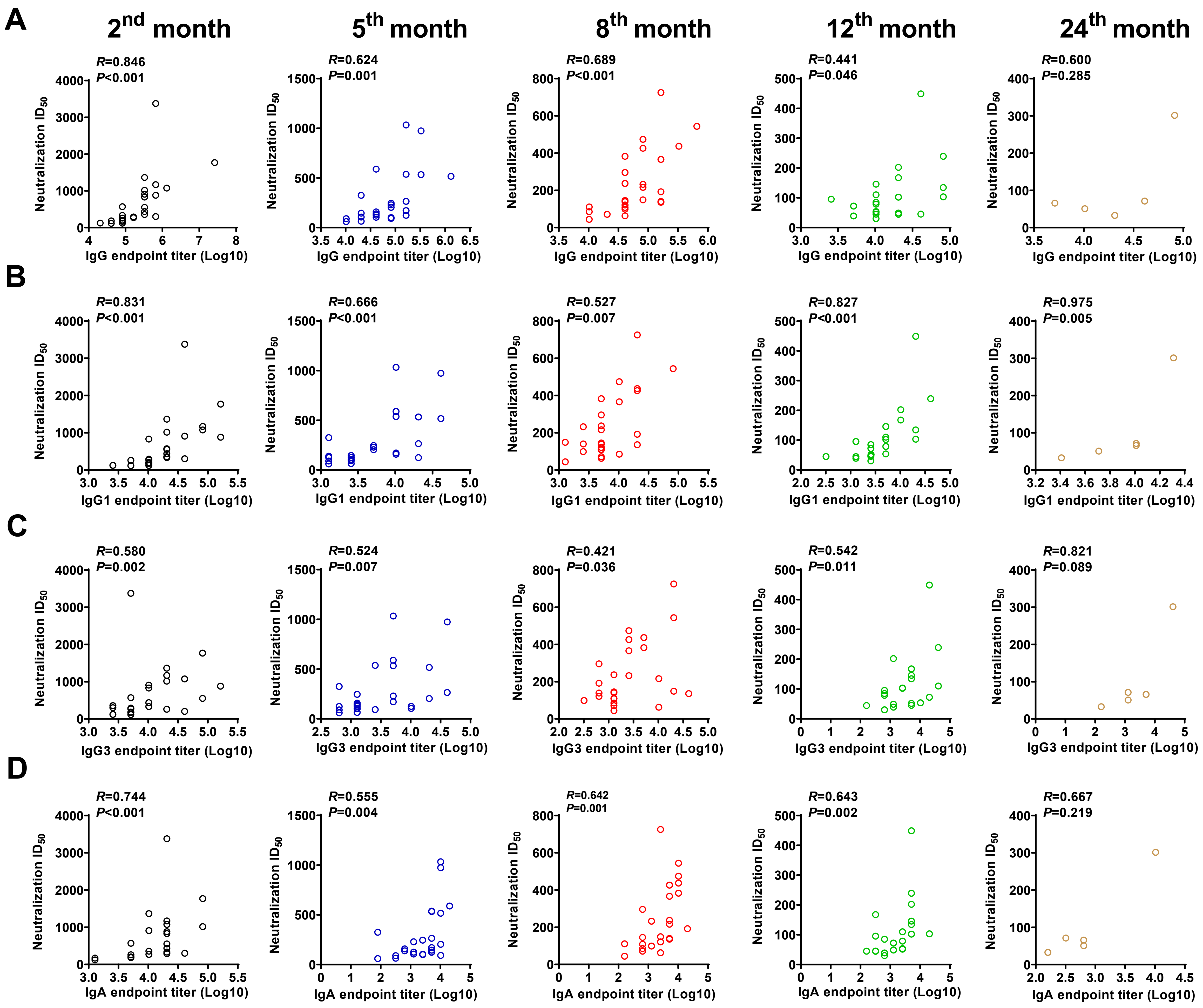

### Supporting Figure 7

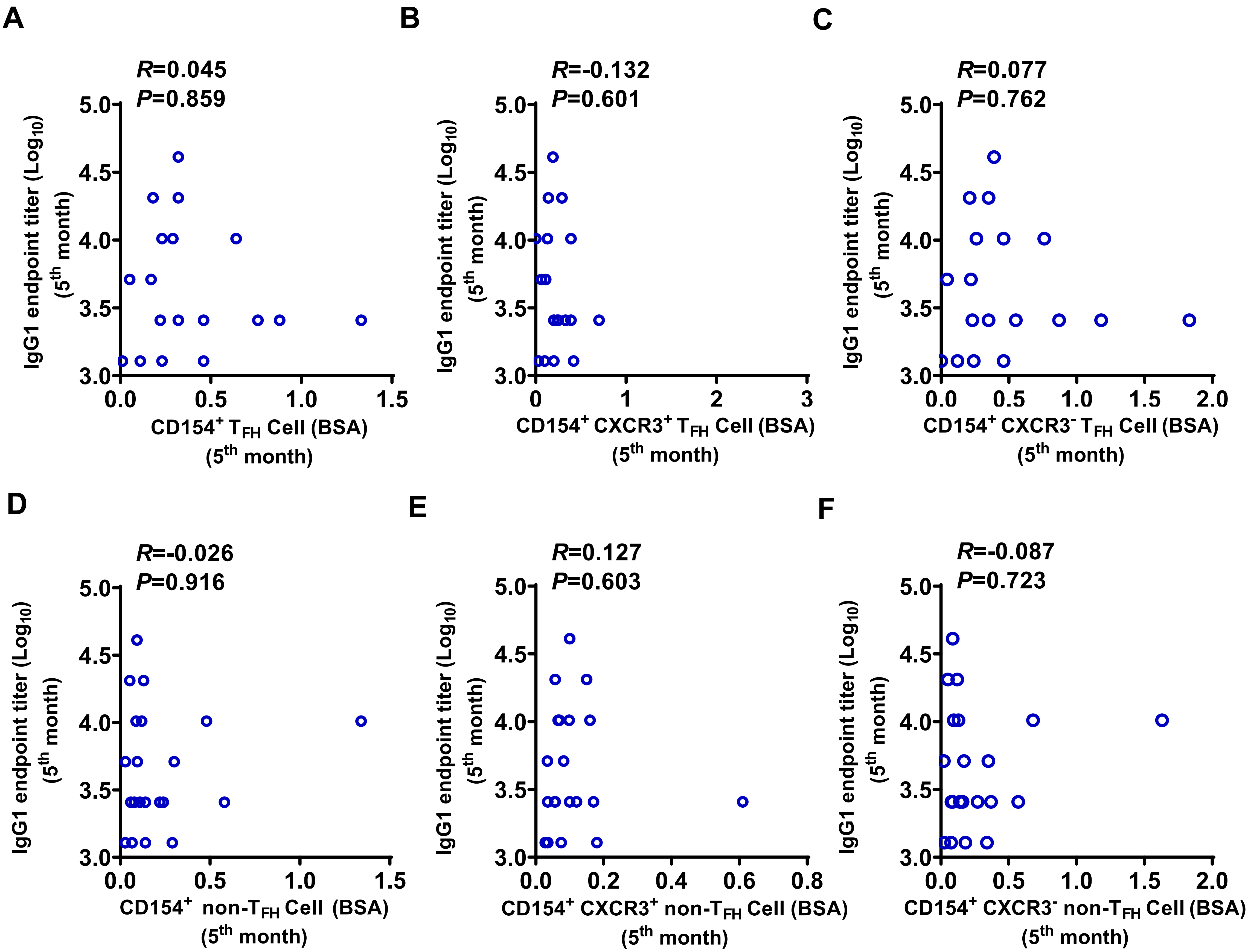

### Supporting Figure 8

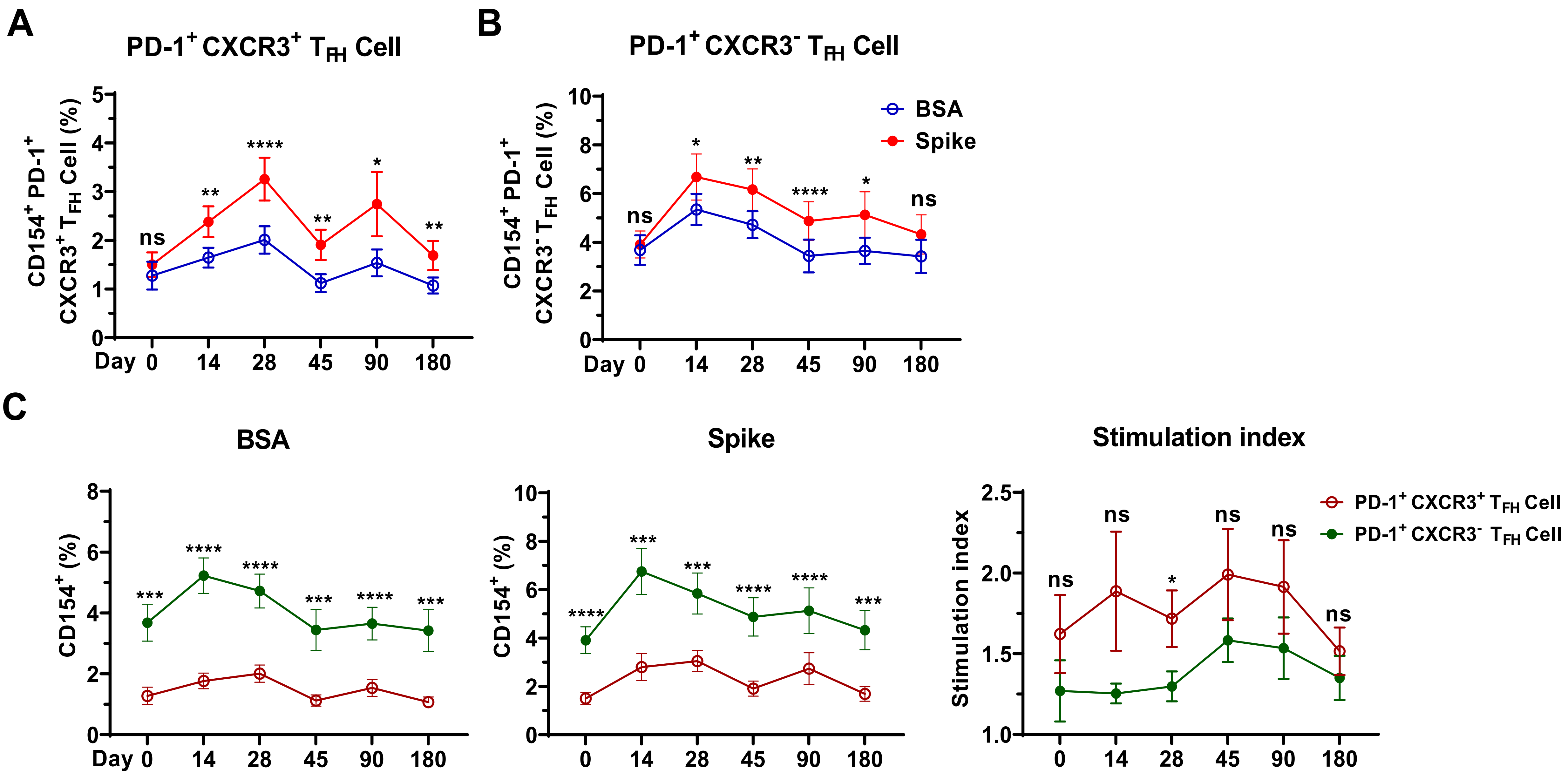

### Supporting Figure 9

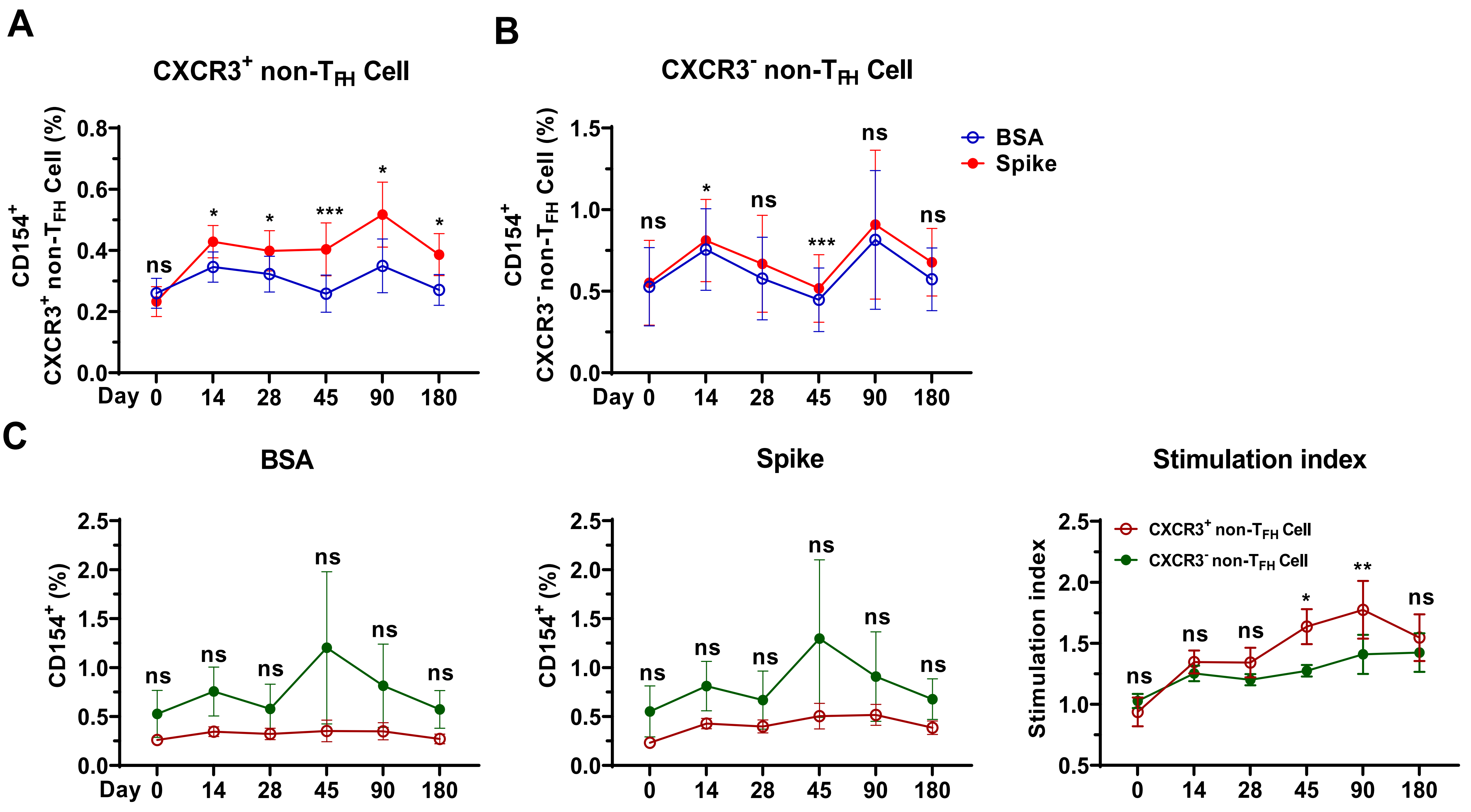

### Supporting Figure 10

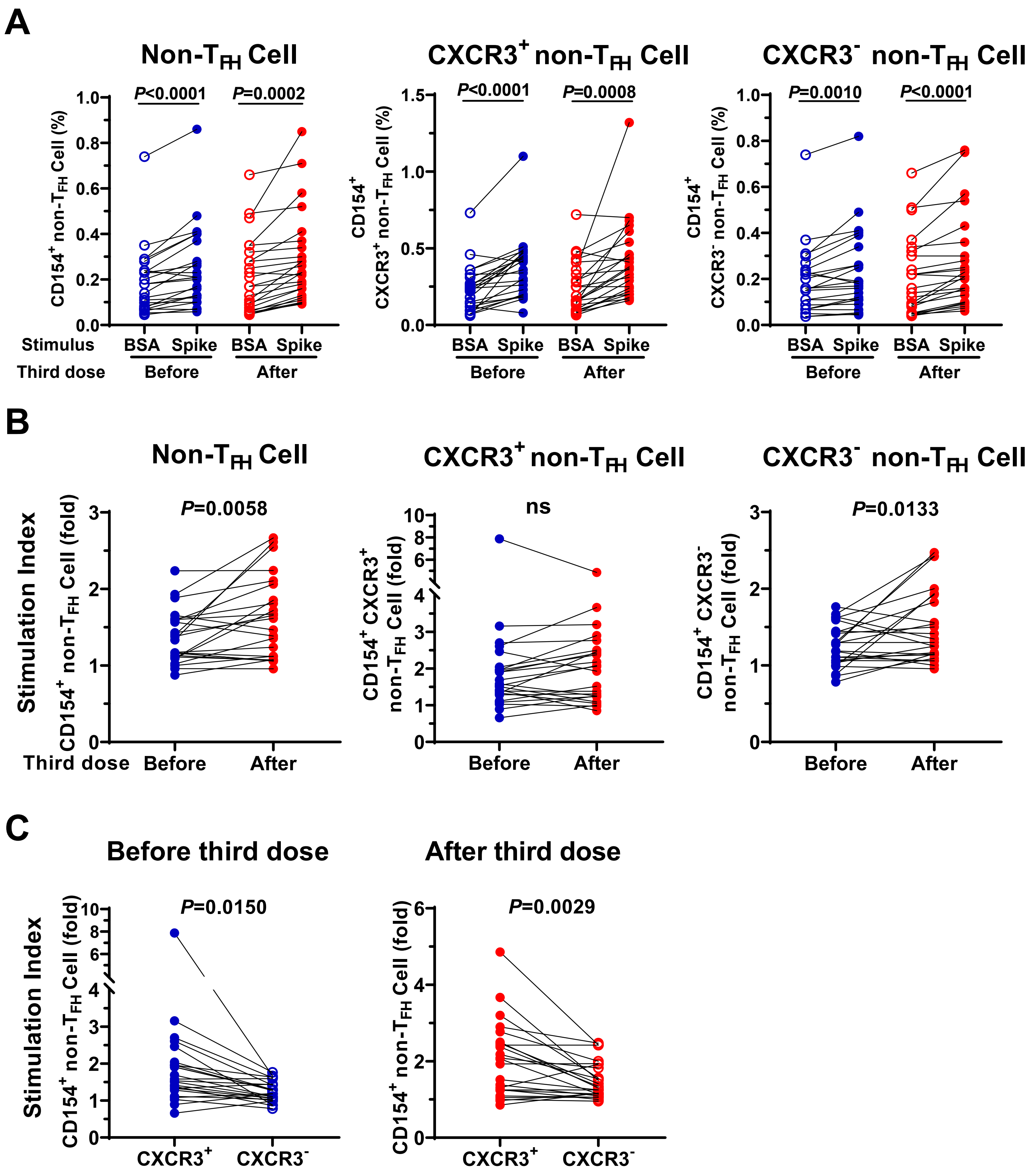

**Supporting Figure 11**

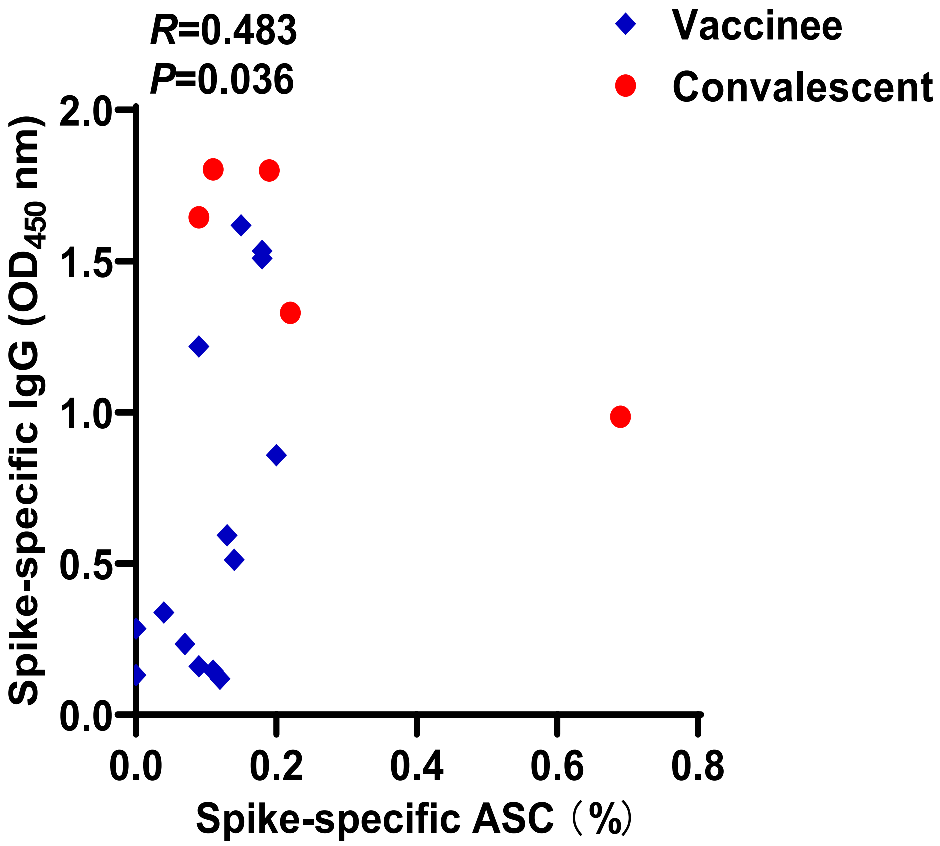
