## Supporting Tables for "Spike-specific CXCR3^+^ T_FH_ cells play a dominant functional role in supporting antibody responses in SARS-CoV-2 infection and vaccination"

### Table 1. Baseline characteristics of participants with COVID-19 recruited in this study

| Patient | Age (years) | Sex | Disease severity | 2nd month | | 5th month | | 8th month | | 12th month | | 24th month | |
|---|---|---|---|---|---|---|---|---|---|---|---|---|---|
| | | | | Days* | PBMCs availability | Days* | PBMCs availability | Days* | PBMCs availability | Days* | PBMCs availability | Days* | PBMCs availability |
| **Patient.1** | 29 | Male | Non severe | 48 | + | 149 | - | 255 | + | 382 | + | NA | |
| **Patient.2** | 48 | Male | Non severe | 42 | + | 143 | - | 249 | + | 376 | - | NA | |
| **Patient.3** | 45 | Female | Non severe | 44 | + | 143 | + | 249 | - | 387 | + | NA | |
| **Patient.4** | 20 | Female | Non severe | 54 | + | 149 | + | 255 | + | NA | | NA | |
| **Patient.5** | 50 | Female | Non severe | 47 | + | 142 | - | 248 | - | 382 | - | NA | |
| **Patient.6** | 69 | Female | Non severe | 52 | - | 179 | + | 248 | + | 375 | + | NA | |
| **Patient.7** | 46 | Female | Non severe | 59 | - | 149 | + | 255 | + | 375 | + | NA | |
| **Patient.8** | 28 | Female | Non severe | 54 | + | 144 | + | 250 | + | NA | | NA | |
| **Patient.9** | 21 | Female | Non severe | 67 | + | 173 | + | 242 | + | NA | | NA | |
| **Patient.10** | 43 | Male | Non severe | 48 | + | 186 | + | 255 | + | 369 | + | NA | |
| **Patient.11** | 23 | Male | Non severe | 62 | - | 182 | + | 251 | - | 382 | + | NA | |
| **Patient.12** | 43 | Female | Non severe | 63 | + | 190 | + | 259 | + | 379 | + | NA | |
| **Patient.13** | 30 | Female | Non severe | 65 | + | 171 | + | 240 | + | NA | | NA | |
| **Patient.14** | 45 | Female | Non severe | 76 | - | 182 | + | 251 | + | 386 | + | NA | |
| **Patient.15** | 22 | Male | Non severe | 66 | + | 172 | + | 241 | + | 367 | + | NA | |
| **Patient.16** | 43 | Male | Non severe | 57 | + | 184 | + | 253 | - | 388 | + | 742 | + |
| **Patient.17** | 47 | Male | Non severe | 62 | + | 145 | + | 251 | - | NA | | NA | |
| **Patient.18** | 50 | Female | Non severe | 58 | + | 153 | + | 259 | + | 368 | + | NA | |
| **Patient.19** | 47 | Female | Non severe | 45 | + | 128 | + | 234 | + | 371 | + | NA | |
| **Patient.20** | 47 | Male | Severe | 53 | + | 154 | + | 260 | + | NA | | NA | |
| **Patient.21** | 64 | Male | Severe | 56 | + | 183 | + | 252 | + | NA | | NA | |

| | | | | | | | | | | | | | |
|---|---|---|---|---|---|---|---|---|---|---|---|---|---|
| **Patient.22** | 72 | Female | Severe | 55 | - | 145 | - | 251 | + | NA | | NA | |
| **Patient.23** | 34 | Female | Severe | 51 | + | 183 | - | 252 | + | 378 | + | NA | |
| **Patient.24** | 67 | Female | Severe | 86 | + | 192 | - | 261 | + | NA | | NA | |
| **Patient.25** | 54 | Male | Severe | 43 | + | 175 | - | 244 | + | 361 | + | NA | |
| **Patient.26** | 52 | Male | Non severe | NA | | NA | | NA | | 370 | + | NA | |
| **Patient.27** | 22 | Female | Non severe | NA | | NA | | NA | | 370 | + | NA | |
| **Patient.28** | 59 | Female | Severe | NA | | NA | | NA | | 381 | + | NA | |
| **Patient.29** | 30 | Male | Non severe | NA | | NA | | NA | | 370 | + | NA | |
| **Patient.30** | 64 | Female | Non severe | NA | | NA | | NA | | 381 | + | NA | |
| **Patient.31** | 18 | Female | Non severe | NA | | NA | | NA | | 367 | - | NA | |
| **Patient.32** | 43 | Male | Non severe | NA | | NA | | NA | | 381 | - | NA | |
| **Patient.33** | 30 | Female | Non severe | NA | | NA | | NA | | 375 | + | NA | |
| **Patient.34** | 43 | Male | Severe | NA | | NA | | NA | | NA | | 731 | + |
| **Patient.35** | 30 | Female | Non severe | NA | | NA | | NA | | NA | | 706 | + |
| **Patient.36** | 52 | Male | Non severe | NA | | NA | | NA | | NA | | 701 | + |
| **Patient.37** | 43 | Female | Non severe | NA | | NA | | NA | | NA | | 701 | + |
| Median | 43 | | | 55.00 | | 171.00 | | 251.00 | | 375.50 | | 706.00 | |
| (IQR) | (30.00-51.00) | | | (48.00-62.50) | | (145.00-182.50) | | (248.00-255.00) | | (370.00-381.75) | | (701.00-736.50) | |

***Specific day that samples were collected for each participant. In the manuscript, the 2nd, 5th, 8th, 12th and 24th month were used for simplicity. +: PBMCs were collected for this experiment; -: PBMCs were not collected; NA: Sample were not available. IQR: Interquartile range.**

**Table 2. Baseline characteristics of vaccine recipients recruited in this study**

| Vaccine recipient | Age (years) | Sex | PBMCs availability | | | | | | | |
|---|---|---|---|---|---|---|---|---|---|---|
| | | | Day0 | Day14 | Day28 | Day45 | Day90 | Day180 | Before Third dose | After Third dose |
| **Vaccine recipient 1** | 37 | Female | - | + | - | - | + | - | NA | NA |
| **Vaccine recipient 2** | 33 | Female | + | - | + | + | + | - | NA | NA |
| **Vaccine recipient 3** | 26 | Female | + | - | + | + | + | + | NA | NA |
| **Vaccine recipient 4** | 26 | Female | + | + | + | + | + | + | NA | NA |
| **Vaccine recipient 5** | 40 | Male | + | + | + | + | + | + | + | + |
| **Vaccine recipient 6** | 30 | Female | + | + | + | + | + | + | NA | NA |
| **Vaccine recipient 7** | 33 | Female | + | + | + | - | + | + | NA | NA |
| **Vaccine recipient 8** | 35 | Female | + | + | + | + | + | + | NA | NA |
| **Vaccine recipient 9** | 28 | Female | + | + | + | + | + | + | + | + |
| **Vaccine recipient 10** | 25 | Female | + | + | + | + | + | + | + | + |
| **Vaccine recipient 11** | 56 | Female | + | + | + | + | + | + | NA | NA |
| **Vaccine recipient 12** | 42 | Male | + | + | + | + | + | - | + | + |
| **Vaccine recipient 13** | 33 | Female | + | + | + | + | + | - | + | + |
| **Vaccine recipient 14** | 31 | Male | + | + | - | + | + | + | + | + |
| **Vaccine recipient 15** | 39 | Female | + | + | + | + | + | + | NA | NA |
| **Vaccine recipient 16** | 36 | Female | + | + | + | + | + | - | + | + |
| **Vaccine recipient 17** | 33 | Female | + | + | + | + | + | - | NA | NA |
| **Vaccine recipient 18** | 51 | Male | + | + | + | + | + | - | + | + |
| **Vaccine recipient 19** | 45 | Male | + | + | + | + | + | + | + | + |
| **Vaccine recipient 20** | 32 | Female | + | + | + | + | + | + | + | + |
| **Vaccine recipient 21** | 45 | Female | + | + | + | + | + | + | + | + |
| **Vaccine recipient 22** | 34 | Female | + | + | - | + | + | + | NA | NA |

| | | | | | | | | | | |
|---|---|---|---|---|---|---|---|---|---|---|
| **Vaccine recipient 23** | 32 | Female | + | + | + | + | + | + | NA | NA |
| **Vaccine recipient 24** | 36 | Female | - | + | + | + | + | + | + | + |
| **Vaccine recipient 25** | 31 | Female | + | + | + | + | + | + | NA | NA |
| **Vaccine recipient 26** | 40 | Female | + | + | + | + | + | + | + | + |
| **Vaccine recipient 27** | 29 | Female | NA | NA | NA | NA | NA | NA | + | + |
| **Vaccine recipient 28** | 48 | Male | NA | NA | NA | NA | NA | NA | + | + |
| **Vaccine recipient 29** | 47 | Male | NA | NA | NA | NA | NA | NA | + | + |
| **Vaccine recipient 30** | 49 | Male | NA | NA | NA | NA | NA | NA | + | + |
| **Vaccine recipient 31** | 31 | Female | NA | NA | NA | NA | NA | NA | + | + |
| **Vaccine recipient 32** | 49 | Female | NA | NA | NA | NA | NA | NA | + | + |
| **Vaccine recipient 33** | 43 | Female | NA | NA | NA | NA | NA | NA | + | + |
| **Vaccine recipient 34** | 38 | Female | NA | NA | NA | NA | NA | NA | + | + |
| **Vaccine recipient 35** | 38 | Female | NA | NA | NA | NA | NA | NA | + | + |
| **Vaccine recipient 36** | 37 | Female | NA | NA | NA | NA | NA | NA | + | + |
| **Vaccine recipient 37** | 28 | Female | NA | NA | NA | NA | NA | NA | + | + |
| **Median** | 36 | | | | | | | | | |
| **(IQR)** | (31-42.5) | | | | | | | | | |

**+: PBMCs were collected for this experiment; -: PBMCs were not collected; NA: Sample were not available. IQR: Interquartile range.**
